# *De novo* macrocyclic peptides for inhibiting, stabilising and probing the function of the Retromer endosomal trafficking complex

**DOI:** 10.1101/2020.12.03.410779

**Authors:** Kai-En Chen, Qian Guo, Yi Cui, Amy K. Kendall, Timothy A. Hill, Ryan J. Hall, Joanna Sacharz, Suzanne J. Norwood, Boyang Xie, Natalya Leneva, Zhe Yang, Rajesh Ghai, David A. Stroud, David Fairlie, Hiroaki Suga, Lauren P. Jackson, Rohan D. Teasdale, Toby Passioura, Brett M. Collins

## Abstract

The Retromer complex (Vps35-Vps26-Vps29) is essential for endosomal membrane trafficking and signalling. Mutations in Retromer cause late-onset Parkinson’s disease, while viral and bacterial pathogens can hijack the complex during cellular infection. To modulate and probe its function we have created a novel series of macrocyclic peptides that bind Retromer with high affinity and specificity. Crystal structures show the majority of cyclic peptides bind to Vps29 via a Pro-Leu-containing sequence, structurally mimicking known interactors such as TBC1D5, and blocking their interaction with Retromer *in vitro* and in cells. By contrast, macrocyclic peptide RT-L4 binds Retromer at the Vps35-Vps26 interface and is a more effective molecular chaperone than reported small molecules, suggesting a new therapeutic avenue for targeting Retromer. Finally, tagged peptides can be used to probe the cellular localisation of Retromer and its functional interactions in cells, providing novel tools for studying Retromer function.

## INTRODUCTION

Endosomal compartments serve as central hubs for transmembrane protein and lipid sorting, and as platforms for cell signalling. Transmembrane protein cargos that arrive in the endosomal network via endocytosis or anterograde trafficking are routed either for lysosomal degradation or recycling to other compartments including the trans-Golgi network (TGN) and the cell surface. Endosomal trafficking thus plays a pivotal role in maintaining cellular homeostasis and is controlled by a number of essential protein machineries ^1–3^.

The evolutionarily conserved Retromer complex is a 150 kDa heterotrimer composed of Vps35, Vps29 and Vps26; with two paralogues Vps26A or Vps26B in vertebrates ^4^. Retromer is a master regulator of endosomal dynamics responsible for cargo sorting and recycling within tubulovesicular transport carriers ^2, 5–9^, and in higher eukaryotes cooperates with an array of cargo adaptors and accessory proteins to allow membrane recruitment, cargo sorting and trafficking to occur ^2, 3, 10^. Known accessory or regulatory proteins include the small GTPase Rab7, the Rab7 GTPase activating protein (GAP) TBC1 domain family member 5 (TBC1D5), VPS9-ankyrin-repeat protein (VARP/Ankrd27), and Fam21, a subunit of the WASP and scar homology (WASH) complex. The known cargo adaptors are derived from the sorting nexin (SNX) protein family and include SNX3 and SNX27. SNX3-Retromer mediated trafficking is primarily thought to mediate the trafficking of cargos from endosomes to the TGN ^11, 12^, whereas SNX27-Retromer is important for retrieval of endocytosed cargos from endosomes back to the cell surface ^13–15^.

Retromer mutation or dysregulation in humans leads to defective endosomal and lysosomal function implicated in neurodegenerative disorders including Parkinson’s disease (PD) ^16–30^, Alzheimer’s disease (AD) ^31–37^ and amyotrophic lateral sclerosis (ALS)^38^. Notably, generalised dysregulation of endosomal, lysosomal and autophagic organelles is a common hallmark of neurodegenerative disorders including familial and sporadic AD, PD, ALS and hereditary spastic paraplegia (HSP) ^20, 39–44^. Retromer dysfunction impacts endosomal and lysosomal homeostasis in neurons and microglia in multiple ways ^20, 45^; involving deficits in regulatory protein interactions such as WASH and Leucine-rich repeat kinase 2 (LRRK2)^19, 23, 27^, mis-sorting of specific endosomal cargos ^18, 26, 28, 46–51^, defects in mitochondrial function and mitophagy ^21, 29, 52, 53^, and other widespread deficiencies in lysosomal and autophagic degradation of toxic material ^23, 24, 32, 34, 37, 54–57^.

Retromer is also a prominent target hijacked by intracellular pathogens to facilitate their transport and replication. This includes viruses such as SARS-CoV-2, human immunodeficiency virus (HIV), hepatitis C virus (HCV), and human papilloma virus (HPV) ^58–70^, and intracellular bacteria such as *Coxiella burnetii* and *Legionella pneumophila* ^71–77^. Mechanistic studies have shown that the secreted effector protein RidL from *L. pneumophila* binds directly to Vps29, competing with endogenous regulators TBC1D5 and VARP, inhibiting Retromer mediated cargo transport, and supporting the growth of *L. pneumophila* in intracellular endosome-derived vacuoles ^71, 73, 76, 77^. Similarly, the minor capsid protein L2 from human Papilloma virus (HPV) is thought to hijack Retromer-SNX3-mediated endosomal transport by mimicking sequence motifs found in endogenous cargoes such as the cation-independent mannose-6-phosphate receptor (CI-MPR) and divalent metal transporter 1-II (DMT1-II) ^63, 66, 78, 79^.

Given the importance of Retromer in endosomal trafficking, neurodegenerative disease, and cellular infection, there is significant interest in developing molecular approaches to either inhibit or enhance Retromer activity. Inhibition of Retromer may provide a novel avenue for targeting infectious pathogens; with peptide-based inhibitors of Retromer, derived from the HPV L2 protein, able to reduce infection by HPV by slowing the normal retrograde transport of incoming virions ^66, 78, 79^. Conversely, because of its neuroprotective role, it has been proposed that a Retromer-binding ‘molecular chaperone’ could be used to enhance Retromer function in diseases including AD and PD ^80–83^, with the goal of boosting normal endosome-dependent clearance pathways to reduce accumulation of toxic aggregates of proteins such as amyloid *β* (A*β*), tau and *α*-synuclein. Previously a small molecule called R55 (a thiophene thiourea derivative, also called TPT-260) was identified, which can bind with modest affinity to Retromer at the interface between Vps35 and Vps29 and stabilise its structure ^84^. This molecule has since been shown to have activity in stabilising Retromer in cells, and as predicted can enhance the transport of essential receptors and reduce the accumulation of toxic material including A*β* and *α*-synuclein in cell, fly, and mouse models ^54, 84–90^. Recently a derivative of R55 was found to improve Retromer stability and lysosomal dysfunction in a model of ALS ^38^. Nonetheless, the low potency and specificity of these compounds mean that other molecules are actively being sought.

We have adopted a novel screening strategy, referred to as the RaPID (Random nonstandard Peptides Integrated Discovery) system, to identify a series of eight *de novo* macrocyclic peptides with high affinity and specificity for Retromer, possessing either inhibitory or stabilising activities. These peptides bind to Retromer with affinities (*K*_d_) ranging from 0.2 to 850 nM and can be classified into two groups based on their binding sites. Most interact specifically with the Vps29 subunit, and crystal structures show that these peptides associate with a highly conserved pocket on Vps29 that is also used by the accessory proteins TBC1D5 and VARP, as well as the bacterial protein RidL, and are potent inhibitors of both TBC1D5 and VARP binding. We also identified one peptide, RT-L4, that shows significant promise as a molecular chaperone. RT-L4 stabilizes Retromer assembly via binding to the interface between Vps26 and Vps35, does not disrupt Retromer’s association with known accessory proteins and cargo adaptors, and indeed allosterically enhances binding to several ligands including SNX27 and TBC1D5. Finally, we show that these macrocyclic peptides can also be used as tools for probing Retromer function. Using reversible cell permeabilization and fluorescent peptides, we demonstrate that they can specifically co-label Vps35-positive endosomal structures and can be used as baits for isolating Retromer from cells. These macrocyclic peptides thus provide novel research tools to enhance our understanding of Retromer-mediated endosomal trafficking and suggest potential new avenues for developing therapeutic modifiers of Retromer function.

## RESULTS

### Identification of highly potent Retromer-binding macrocyclic peptides

The procedure of the RaPID system (**Fig. 1A**) exploits the diverse molecular topology of macrocyclic peptide populations numbering >10^12^ unique sequences to enrich for and amplify low abundance, high-affinity ligands ^27, 91–94^. We performed the RaPID selection using purified His-tagged human Retromer complex as bait: a puromycin-linked semi-randomized messenger RNA library, of the general form AUG-(NNK)_4-15_-UGC, was translated in an *in vitro* translation reaction to generate peptides covalently linked to their cognate mRNAs through the puromycin moiety. Macrocyclization was effected through genetic code reprogramming of initiating AUG (methionine) codons to incorporate either *N*-chloroacetyl-L-tyrosine (ClAc-L-Tyr) or *N*-chloroacetyl-D-tyrosine (ClAc-D-Tyr), leading to spontaneous reaction with a downstream UGC-encoded cysteine to form a library of thioether bridged cyclic peptides each linked to their cognate mRNA (total diversity >10^12^ molecules). Retromer ligands were then identified through iterative cycles of affinity selection against immobilised Retromer complex followed by RT-PCR recovery, transcription and regeneration of peptide-mRNA fusion libraries, with deconvolution of the final enriched library achieved through next-generation sequencing.

**Figure 1.**
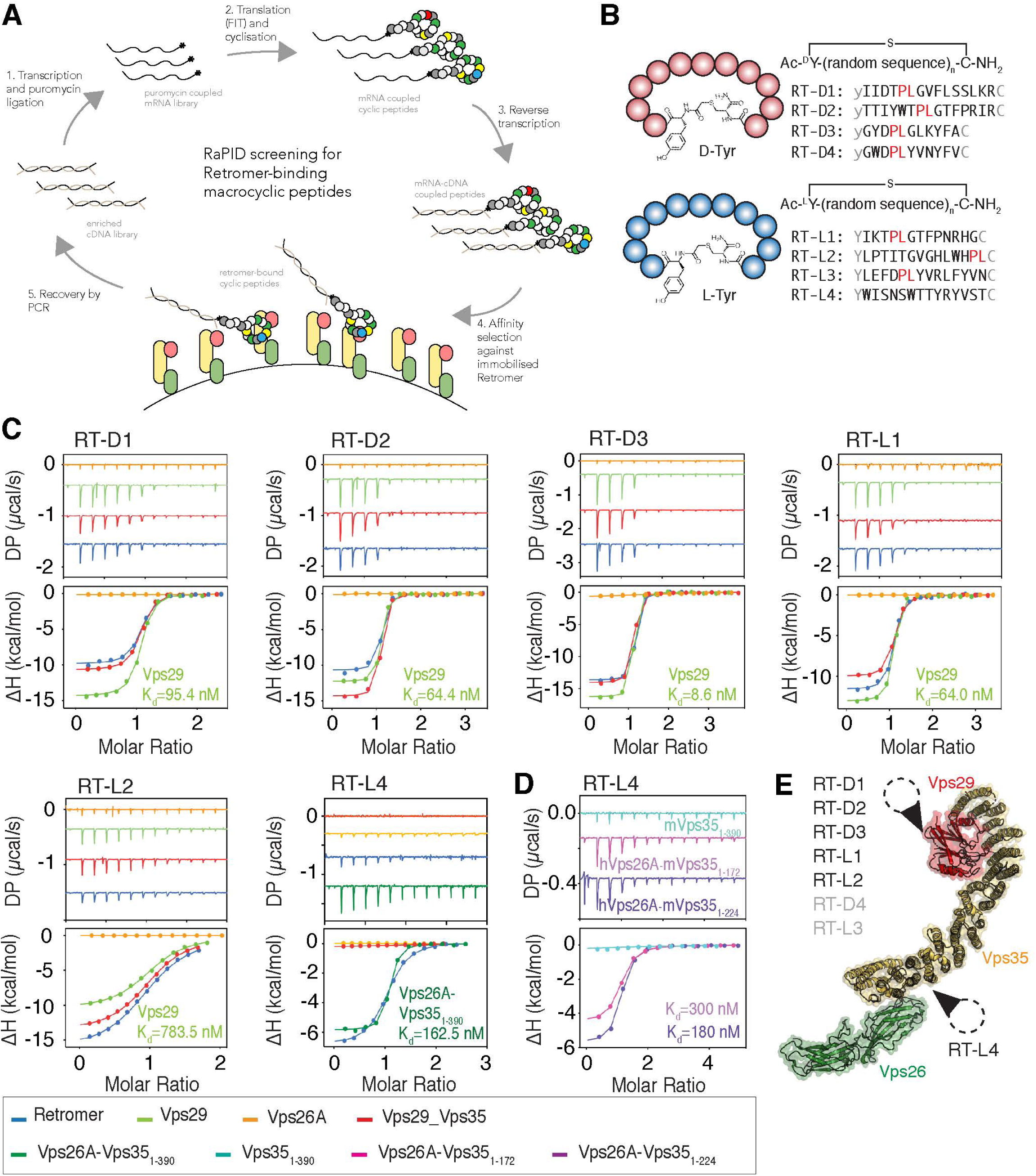
Cyclic peptides reveal strong binding characteristics with Retromer. (**A**) Schematic diagram showing the RaPID system used to screen for cyclic peptides binding to Retromer. (**B**) Eight Retromer-binding macrocyclic peptides were identified with either N-chloroacetyl-_D_-tyrosine or N-chloroacetyl-_L_-tyrosine as initiating residues. (**C**) Binding of RT-D1, RT-D2, RT-D3, RT-L1, RT-L2, and RT-L4 with Retromer (blue), Vps29 (light green), Vps26A (orange), Vps29 – Vps35 (red) and Vps26A – Vps35_1-390_ (dark green) by ITC. SPR binding curves for each peptide are shown in **Fig. S3**. (**D**) ITC thermogram showing that addition of α-helix 8 and 9 of Vps35 (residues 173 to 224) contributes to RT-L4 binding affinity but is not essential for interaction. All ITC graphs represent the integrated and normalized data fit with 1 to 1 ratio binding. The binding affinity (*K*_d_) is given as mean of at least three independent experiments (**Table S2**). (**E**) Relative binding position of each cyclic peptide to Retromer based on the ITC measurements indicated on the structure of the mouse Retromer complex (PDB ID 6VAC) ^100^.

We selected the four most abundant peptides from each of the ClAc-L-Tyr or ClAc-D-Tyr initiated libraries (**Fig. 1B**; **Fig. S1; Table S1**) for further characterisation, and each was synthesised using standard Fmoc chemistry. These were validated against the Retromer complex by surface plasmon resonance (SPR) and all were confirmed to bind very strongly to Retromer with affinities (*K*_d_s) in the range of <0.2 to 30 nM (**Fig. S2**). Secondary SPR comparisons showed that these molecules also bound the purified Retromer complex from *Danio rerio* (zRetromer) (**Fig. S2**) and the thermophilic yeast *Chaetomium thermophilum* (ctRetromer) with a similar range of binding affinities, indicating that they associate with evolutionarily conserved sites in the complex.

Macrocyclic peptide binding to Retromer was validated using isothermal titration calorimetry (ITC). Among the eight cyclic peptides tested, six were confirmed to bind Retromer with nanomolar binding affinity while the peptides RT-D4 and RT-L3 were not sufficiently soluble for ITC experiments (**Figs. 1C and 1D; Table S2**). Affinities for Retromer measured by ITC were systematically lower but correlated well with those measured by SPR, ranging from 25 to 850 nM. These affinities are comparable to or better than the high affinities for the Retromer binding regulatory protein TBC1D5 (220 nM) ^95^, or the bacterial effector RidL (150 nM) ^76, 77^.

The Retromer structure consists of a long Vps35 *α*-helical solenoid, with Vps26A or Vps26B bound to the N-terminus and Vps29 bound at the C-terminus (**Fig. 1E**) ^96–100^. We examined the specific subunits of Retromer required for binding each of the cyclic peptides by testing either Vps29 alone, Vps26A alone, the Vps35-Vps29 heterodimer, or sub-complexes of Vps26A with N-terminal fragments of Vps35 (**Fig. 1D; Table 1)**. Full-length Vps35 is relatively unstable on its own and was not tested separately. Interestingly, RT-D1, RT-D2, RT-D3, RT-L1 and RT-L2 all bind specifically to either Vps29 alone or to the Vps35-Vps29 heterodimer, but not to Vps26A. Their affinities for Vps29 alone were not significantly different to their binding to the Retromer holo-complex or Vps35-Vps29 heterodimer, ranging from 8 to 783 nM. This indicates they bind specifically and exclusively to the Vps29 subunit. Although peptides RT-D4 and RT-L3 were not tested for binding specific subunits, it is likely they also bind to Vps29 as they possess a Pro-Leu motif that we show below mediates Vps29 interaction by the other five peptides (**Figs. 1B and 1E**).

In contrast to other peptides, RT-L4 did not bind to any of the Retromer subunits individually or to the Vps35-Vps29 dimer, but rather only to sub-complexes that contained *both* Vps26A and Vps35 (**Fig. 1C**). N-terminal fragments of Vps35 in complex with Vps26A, including Vps35_1-390_, and Vps35_1-224_ supported binding to the RT-L4 peptide with a similar affinity to the Retromer trimeric holo-complex, while Vps35_1-390_ on its own did not (**Fig. 1D**). When Vps35 was truncated to the shortest region still capable of Vps26A interaction (Vps35_1-172_) a small but consistent decrease in binding affinity was observed, suggesting that *α*-helix 8 in Vps35 (residues 175 to 195) of Vps35 contributes to the binding but is not essential. Overall, we have found that the cyclic peptides RT-D1, RT-D2, RT-D3, RT-L1 and RT-L2 (and likely RT-D4 and RT-L3) bind specifically to Vps29, but RT-L4 binding occurs at the interface between the Vps26A and Vps35 subunits (**Fig. 1E**).

### Macrocyclic peptides bind to Vps29 through a conserved site mimicking endogenous accessory proteins

To understand the molecular basis of how the discovered macrocyclic peptides bind to Retromer, we next co-crystallized Vps29 with RT-D1, RT-D2, RT-D3, RT-L1 and RT-L2. With extensive crystallization trials, we successfully solved structures of human Vps29 in complex with RT-D1, RT-D2, RT-L1 and RT-L2 (**Fig. 2; Table S3**). For RT-D3, the complex structure was successfully determined using Vps29 from *C. thermophilum* (ctVps29). In all five complex structures the conformation of Vps29 is highly similar to apo Vps29 (root mean square deviation ranges from 0.6 Å to 1.3 Å), with cyclic peptide density clearly visible (**Fig. 2A; Fig. S3A**). Surprisingly, although the precise details differed for each macrocyclic peptide, they all bound to the same hydrophobic surface groove on Vps29 composed of multiple conserved residues including Leu2, Leu25, Leu152, Tyr163, Tyr165 and Val174 (human numbering) (**Fig. 2B**). This hydrophobic cavity is highly conserved throughout evolution (**Fig. 2B**) and is located on the opposite surface of Vps29 relative to the Vps35 binding region (**Fig. 2C**).

**Figure 2.**
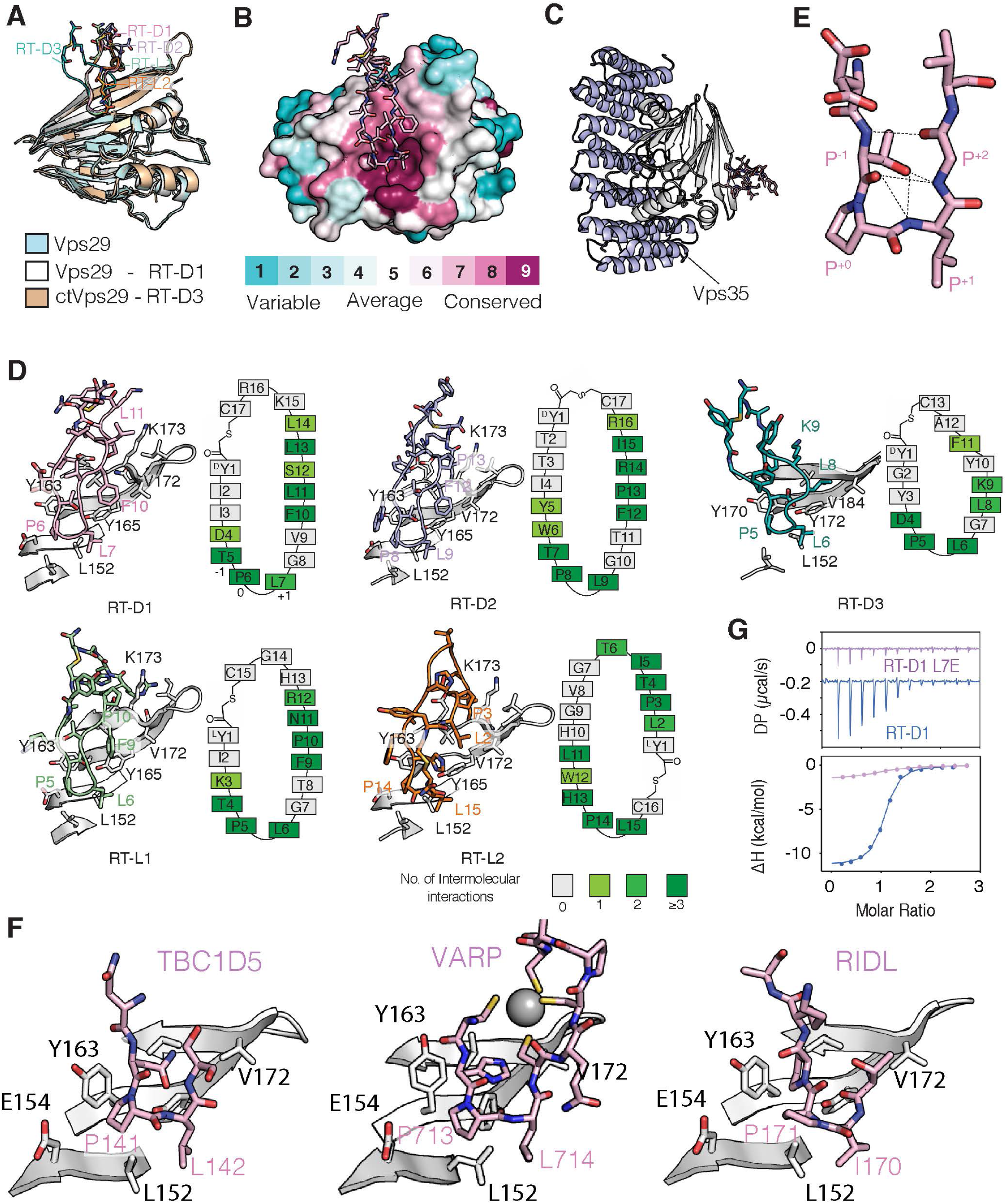
Crystal structures of Vps29 in complex with cyclic peptides. (**A**) Superimposition of the crystal structures of Vps29 in complex with five macrocyclic peptides. (**B**) Sequence conservation mapped onto the hVPS29 structure surface highlights the conserved binding site for each of the cyclic peptides, with RT-D1 shown as an example. (**C**) Superimposition of the Vps29 – RT-D1 peptide complex with the Vps29 - Vps35_483-780_ crystal structure (PDB ID: 2R17) ^96^. The cyclic peptides bind opposite to the Vps35 interface. (D) Details of macrocyclic peptides in stick representation bound to Vps29. Schematic diagrams indicate the number of intermolecular contacts (salt bridge, hydrogen bonds and hydrophobic interactions) of each residue on the peptide with Vps29. The residues on the peptide with 0, 1, 2 and ≥3 contacts are shown in grey, light green, green and dark green boxes respectively. Each peptide utilises a core Pro-Leu sequence forming a *b*-hairpin, labelled as position 0 and 1 for reference. Electron density for each peptide is shown in **Fig. S3**. (E) Close-up of the *β*-hairpin conformation of the bound RT-D1 peptide. (**F**) TBC1D5 (PDB ID 5GTU) ^95^, VARP (PDB ID 6TL0) ^103^ and RIDL (PDB ID 5OSH) ^76^ bind to the same site of Vps29 using a similar *β*-hairpin comprised of a Pro-Leu dipeptide sequence. For clarity, only the key residues involve in the contact are shown. (**G**) ITC thermogram for the titration of RT-D1 (purple) and RT-D1 L7E (blue) with Retromer showing the importance of Pro-Leu motif in the interaction. The graph represents the integrated and normalized data fit with a 1 to 1 binding ratio.

Each cyclic peptide adopts a different conformation when bound to Vps29, as expected from their lack of sequence identity (**Fig. 2D; Fig. S3B**). However, a notable feature of all of the Vps29 binding peptides is a Pro-Leu di-peptide motif present in a *β*-hairpin structure that inserts into the conserved hydrophobic site on Vps29 (**Fig. 1B; Figs. 2D and 2E**). Further analysis reveals the residue in front of the Pro-Leu motif (position -1, with the Pro designated as position ‘0’) is important for stabilizing the β-hairpin like configuration, forming multiple hydrogen bonds with nearby residues including the side-chain hydroxyl group of Vps29 Tyr165 (**Figs. 2D and 2E**). Notably TBC1D5, VARP and the bacterial hijacking molecule RidL all engage Vps29 at the same site, employing a similar Pro-Leu dipeptide at their core (**Fig. 2F**) ^71, 76, 77, 95, 101–103^. The Vps29-binding peptides that we have identified thus mimic these natural interactions but exhibit higher affinities.

In addition to the core Pro-Leu di-peptide motif, the residues at positions +4 to +7 of the cyclic peptides also form extensive contacts with a surface groove composed of Vps29 side chains Val172, Lys173 and Val174 (**Fig. 2D; Fig. S3B**). This particular interaction network is shifted by 8 Å towards Lys188 in the ctVps29 – RT-D3 structure (equivalent to Arg176 in human Vps29), which may explain the weaker binding of ctVps29 to RT-D3 compared to Vps29 (**Fig. 2D; Fig. S3C**). To confirm that the Pro-Leu motif is a key feature for these cyclic peptides to recognize Vps29, we altered the Leu at position +1 in RT-D1 to Glu, binding to Retromer was almost abolished (**Fig. 2G**). Another noteworthy observation was found in the Vps29 – RT-L1 structure, where a potential secondary cyclic peptide binding pocket was identified, surrounded by helix 3 and Ile91 of Vps29 (**Fig. S3D**), a region known to be required for Vps35 binding. In this binding pocket, we found two RT-L1 peptides are bound to each other, forming extensive contacts with residues located on helix 3 and adjacent loop region of Vps29 (**Fig. S3E**). This binding of RT-L1 to a secondary site in Vps29 is likely due to a crystallisation-induced contact, but it could suggest a potential site for targeting the Vps29-Vps35 interaction in the future.

### The RT-L4 macrocyclic peptide is a molecular chaperone that binds to the interface between Vps26A and Vps35

One of our initial goals was to identify potential molecular chaperones that can stabilise the Retromer complex, hence we next assessed the impact of the macrocyclic peptides on the thermal stability of Retromer using differential scanning fluorimetry (DSF) (**Fig. S4A**). The Vps29-binding peptides RT-D1, RT-D2, RT-D3, RT-L1 and RT-L2 all increased the melting temperature (T_m_) of Vps29 upon addition of a 20-fold molar excess (**Fig. S4A**), consistent with their high affinities. In contrast however, none of the Vps29-specific macrocyclic peptides had a significant impact on the thermal stability of the trimeric Retromer holo-complex (**Fig. S4B**). This suggests that increasing the thermal stability of Vps29 alone is insufficient to enhance the stability of the entire Retromer complex in solution. In contrast, the addition of RT-L4 resulted in a substantial 2°C to 6.5°C enhancement in the T_m_ of the Retromer holo-complex in a dose-dependent manner (**Fig. 3A; Figs. S4C and S4D**). Similarly, the RT-L4 peptide resulted in an 8°C increase in T_m_ for the Vps26A-Vps35_1-390_ sub-complex, and as expected in control experiments it did not affect the thermal stability of either Vps26A or Vps29 alone (**Fig. S4C**). Using mass photometry, we found that the Retromer holo-complex was maintained at its trimeric state upon the addition of cyclic peptides, suggesting that the enhancement in the T_m_ of the Retromer was not the result of high-order oligomer formation (**Fig. S5A**). These results together indicate that RT-L4 stabilizes Retromer through its interaction at the interface between Vps26A and Vps35, possibly by acting as a molecular ‘staple’ between the two subunits. In parallel, we also compared the thermal stability effect of RT-L4 against the previously published Retromer chaperone R55 ^84^. Surprisingly we were unable to detect any improvement in the T_m_ of Retromer in the presence of R55 under the same experimental conditions as RT-L4 (**Fig. S5B**). Even in the presence of 1 mM R55 (> 300-fold molar excess) no impact on Retromer stability was detectable (**Fig. S5C**), although a second unfolding event was observed that might indicate partial stabilisation of the Vps29-Vps35 complex ^84^. This difference is likely explained by the much lower binding affinity of R55 to Retromer with a *K*_d_ of 15 μM (**Fig. S5D**). Our results indicate that RT-L4 can act as a potent molecular stabiliser of Retromer in solution.

**Figure 3.**
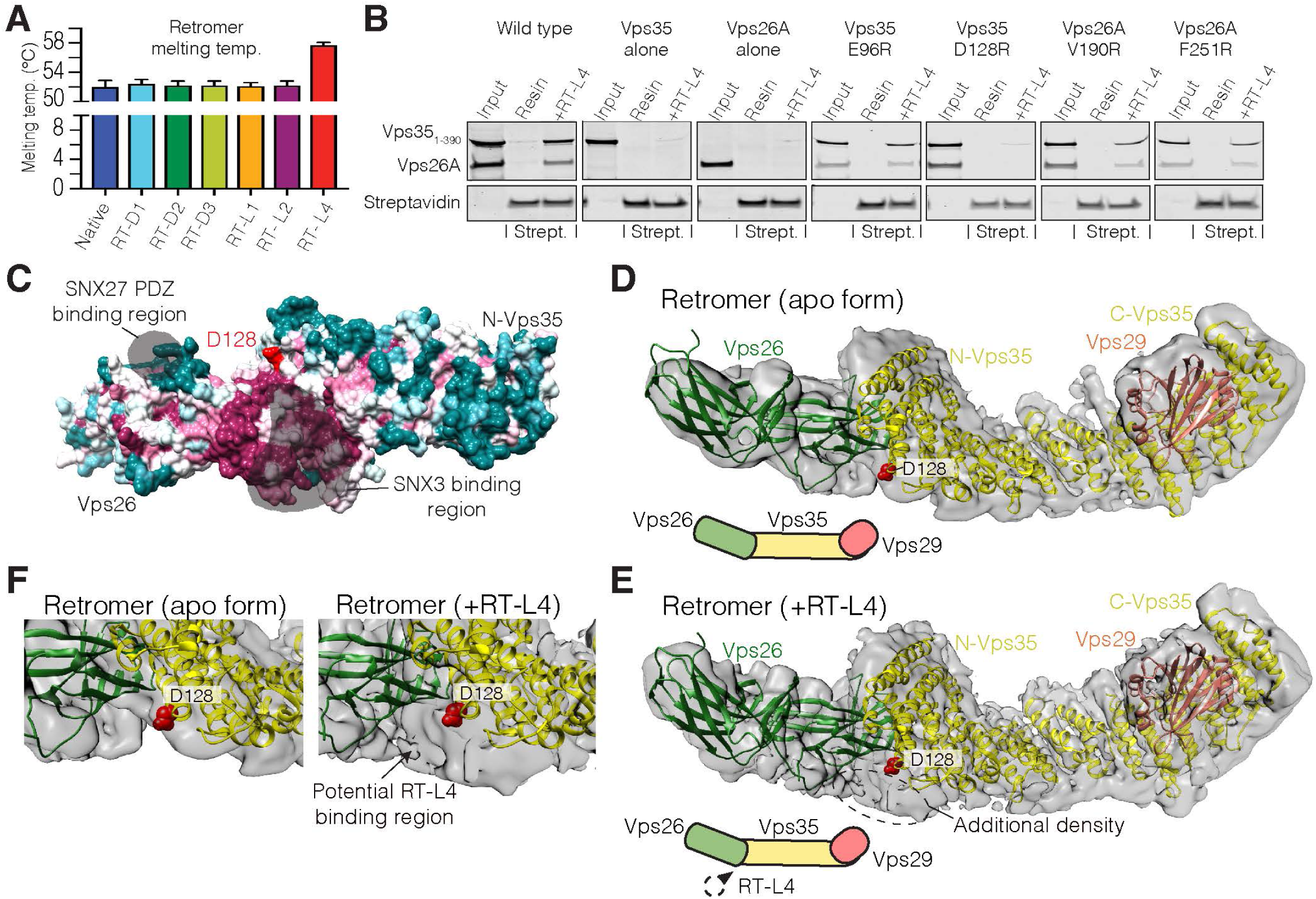
The RT-L4 macrocylic peptide is a molecular chaperone that binds Retromer at the Vps35-Vps26 interface. (**A**) Bar graph summarizing the measured thermal unfolding temperatures (T_m_) of Retromer in the presence of cyclic peptides. Raw data is shown in **Fig. S4**. **(B**) Pull-down assay showing the interaction of biotinylated RT-L4 bound to streptavidin-coated agarose beads with either wild-type Vps26A-Vps35_1-390_ subcomplex or indicated point mutants. Individual Vps26A and Vps35_1-390_ proteins do not bind the peptide, while the D128R mutation in Vps35_1-390_ specifically blocks sub-complex interaction. (**C**) Sequence conservation map of Vps26A-Vps35_12-470_ subcomplex (PDB ID 5F0L) highlighting the proposed RT-L4 binding site, and its relationship to known binding sites for SNX3 ^98^ and SNX27 ^105^. (**D**) CryoEM reconstruction of the human Retromer (apo form) and (**E**) in complex with RT-L4. CryoEM density shown as transparent molecular envelope, with crystal structures of Retromer subcomplexes (PDB ID 2R17 and 5F0L) overlapped to the map (contoured at 4.7σ). The additional density seen on addition of the cyclic peptide added Retromer supports the mutagenesis data indicating RT-L4 binds at the Vps26 and Vps35 interface. (**F**) Enlarged view of the Vps26 and Vps35 interface highlighting the additional density in the CryoEM map of RT-L4 added Retromer but not in the apo form. For clarity, D128 of Vps35 is highlighted in red.

Next, we sought to map the exact binding region of RT-L4 to better investigate the mechanism of its interaction and how this results in the enhanced thermal-stability of Retromer. Using ITC we found that RT-L4 binds to zRetromer with a *K*_d_ of 80 nM, which is highly comparable to human Retromer or the Vps26A-Vps35_1-390_ sub-complex (**Table S2**; **Fig. S6A**). In contrast, RT-L4 binds with lower affinity to ctRetromer with a *K*_d_ of 6 μM (**Table S2**; **Fig. S6B**). As our data indicated that RT-L4 binds to a site that is formed by a combined interface on Vps35-Vps26A, to investigate the residues required to the binding we performed directed mutagenesis of residues surrounding the human Vps35-Vps26A interface based on the previous crystal structure ^98^. Streptavidin agarose beads coated with biotinylated RT-L4 were used to pulldown purified Vps26A-Vps35_1-390_ incorporating several specific point mutations, and this revealed that peptide binding was abolished by the D128R mutation in Vps35 (**Fig. 3B**). The Asp128 side-chain is part of a conserved surface adjacent to the Vps35-Vps26A interface (**Fig. 3C**), and makes a minor contact with the extended N-terminus of SNX3 when in a ternary complex with the SNX3 adaptor and *ΩΦ*[LV] sequence containing-cargo peptides (where *Ω* and *Φ* are aromatic and hydrophobic side-chains) ^3, 98^.

We further explored validated the site where RT-L4 binds Retromer using single particle cryo-electron microscopy (cryoEM). In previous work, wild-type Retromer was found to form a mixed population of hetetrotrimers and multiple higher order oligomers in vitreous ice, so we used an established mutant called “3KE Retromer” (point mutations E615A/D616A/E617A in Vps35) that favours the heterotrimeric species over higher order oligomers for our analyses ^100^. We determined structures of both apo and RT-L4-bound 3KE Retromer under the same conditions to ascertain whether the RT-L4 binding site could be identified (**Fig. 3D-F**; **Figs. S6C-F**; **Table S4**). At the current resolution we detected no structural differences between wild-type heterotrimer ^100^ and the 3KE mutant. Comparison of our reconstructions indicates there is additional density at the Vps26-Vps35 interface in the presence of the RT-L4 peptide, although the estimated resolution of apo and RT-L4/Retromer structures (∼5.0 Å; 0.143 FSC cut-off in RELION) is too low to unambiguously assign peptide density (**Figs. 3D-F**; **Figs. S6C and S6D**). We note RT-L4/Retromer exhibits more preferred orientation in vitreous ice than does apo Retromer, which limits particle views (**Fig. S6D**). We observe “top-down” views of RT-L4/Retromer, but we lack views rotated by 180 degrees (**Fig. S6D**). One explanation for this difference may be relatively low solubility for the RT-L4 peptide, which in turn may influence how RT-L4/Retromer behaves at the air-water interface. Although not conclusive, these reconstructions support our biophysical and mutagenesis data, providing additional evidence RT-L4 binds and stabilizes Retromer at the Vps26/Vps35 interface, but higher resolution data will be required to identify residues in both Retromer and the peptide that specifically mediate the interaction.

### The impact of cyclic peptides on the interactions between Retromer and known regulatory proteins

Given the high affinity and specificity of the macrocyclic peptides for Retromer, it was critical to assess their potential effects on Retromer’s interaction with essential regulatory, accessory and adaptor proteins. We used RT-L4, with its distinct affinity for the Vps26-Vps35 complex, and RT-D3 as a representative Vps29-binding peptide, and tested Retromer binding proteins for which binding mechanisms were known including TBC1D5^95^, SNX3^98^, Fam21^104^, and SNX27^105^ (**Figs. 4A-E**). GST-tagged TBC domain of human TBC1D5 (TBC1D5_TBC_) bound Retromer in pulldowns, and this interaction was inhibited by addition of RT-D3 as expected based on their overlapping binding site (**Fig. 4A**). A similar result was observed using ITC, where we observed binding between TBC1D5_TBC_ and Retromer with a *K*_d_ of 370 nM (consistent with previous reports ^95^), while binding was undetectable in the presence of competing RT-D3 (**Fig. 4B, Table S5**). In contrast, while the addition of RT-D3 blocked Retromer from interacting with TBC1D5_TBC_ as would be predicted from their overlapping binding sites, it had no impact on Retromer binding to either SNX27 or SNX3 as assessed in pulldowns and ITC experiments (**Figs. 4A, 4C, and 4D**). Note that for ITC and pull-down experiments with SNX3, we found it was necessary to include a synthetic peptide corresponding to DMT1-II_550-568_ containing a *ΩΦ*[LV] cargo motif (where *Ω* = an aromatic side-chain and *Φ* = a hydrophobic side-chain) ^98^. In the absence of this peptide we could not detect binding of SNX3 to Retromer, likely because SNX3 and cargo motifs form a co-dependent binding interface with the Retromer complex ^3, 98^. To our surprise, the addition of RT-D3 modestly affected Retromer binding to Fam21 C-terminal LFa repeats 19 to 21 (**Figs. 4A and 4E**), a sequence known to associate with a C-terminal region of Vps35 ^19, 104^. It is therefore likely that Vps29 contributes partially to the binding of LFa sequences of Fam21, and RT-D3 either directly competes with Fam21 or allosterically affects the Fam21 binding site. These results were further validated using qualitative pulldowns from HeLa cell lysates with biotinylated RT-D3 peptide, where specific loss of TBC1D5 and VARP binding was seen, without any effect on the association of other Retromer ligands SNX27, Fam21, SNX3 and SNX27 (**Fig. 4F**). Altogether our data indicates RT-D3 can specifically inhibit binding to TBC1D5, and most likely will also compete with other proteins such as VARP and RidL that bind to the same conserved hydrophobic cavity on Vps29.

**Figure 4.**
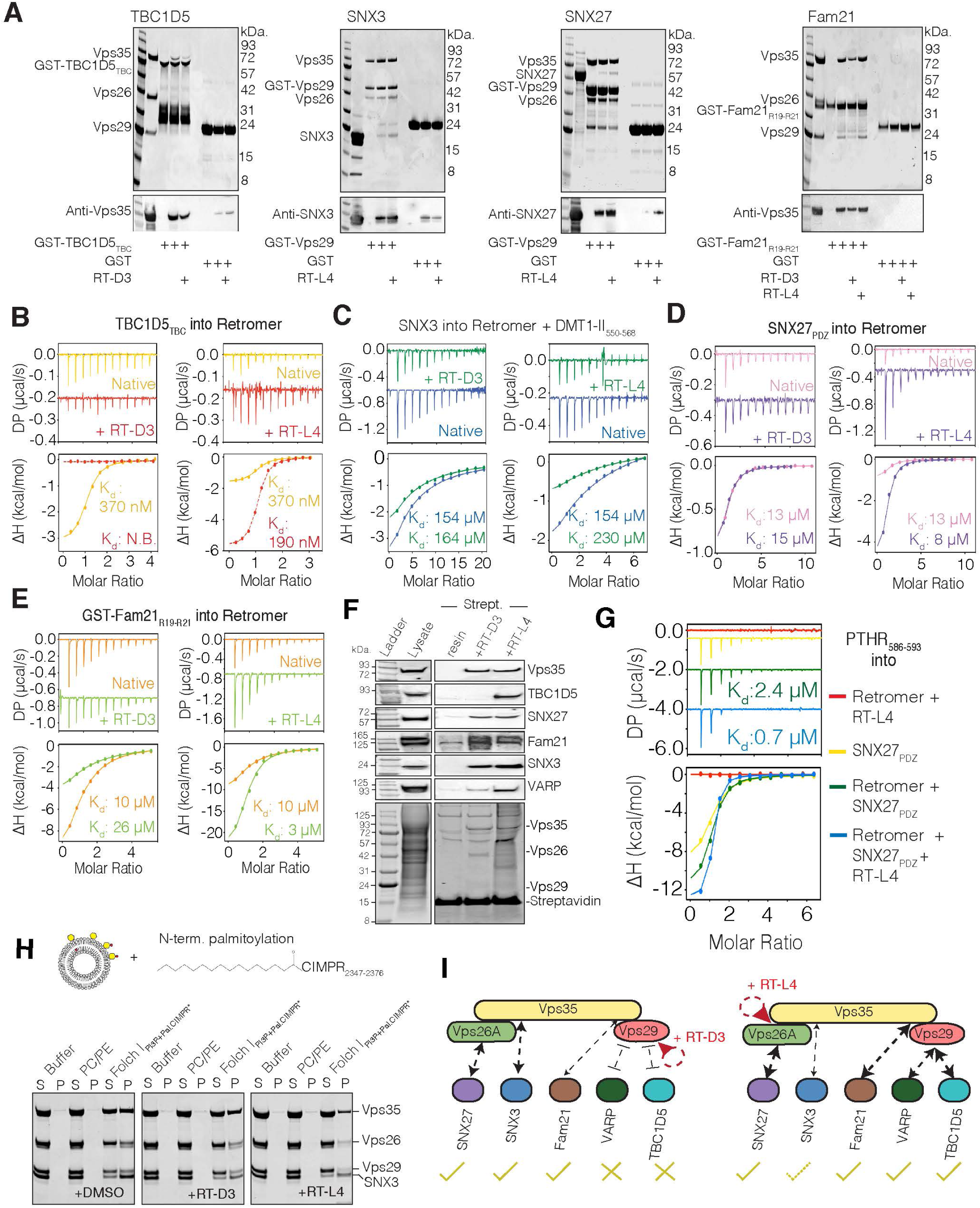
The effect of cyclic peptides on the interaction of Retromer with known regulatory and adaptor proteins. (**A**) Interactions of Retromer with TBC1D5, SNX3, SNX27 and Fam21 in the presence of either RT-D3 or RT-L4. GST-TBC1D5_TBC_ and GST-Fam21_R19-R21_ were used as baits for Retromer, while GST-tagged Retromer (Vps29 subunit) was used as bait for SNX3 and SNX27. (**B**) ITC measurement of TBC1D5_TBC_, (**C**) SNX3 (with DMT1-II_550-568_ present), (**D**) SNX27, and (**E**) GST-Fam21_R19-R21_ with Retromer in the presence or absence of RT-D3 or RT-L4. (**F**) Hela cell lysates were incubated with streptavidin agarose coated with biotinylated RT-D3 or RT-L4 and bound proteins subjected to SDS-PAGE and western blotting with antibodies against indicated proteins. (**G**) ITC measurement of PTHR_586-593_ cargo peptide ^14^ with SNX27_PDZ_ alone (yellow), Retromer + RT-L4 (red), Retromer + SNX27_PDZ_ (green) and Retromer + SNX27_PDZ_ + RT-L4 (blue). RT-L4 allosterically enhances the affinity of Retromer + SNX27_PDZ_ for cargo. The cargo peptide binds to SNX27 All ITC graphs represent the integrated and normalized data fit with a 1 to 1 binding ratio. The binding affinity (K_d_) is given as mean of at least two independent experiments. (**H**) Liposome-binding assay of Retromer with membrane-associated SNX3 cargo complex in the presence of cyclic peptides. Multilamellar vesicles were composed of either control PC/PE lipids, or Folch I lipids containing added PtdIns(3)*P* and N-terminal palmitoylated CIMPR_2347-2376_ peptide (schematic diagram on top). “S” and “P” indicates unbound supernatant and bound pellet respectively. Control experiments are shown in **Fig. S7B** and **S7C**. (**I**) Schematic summarizing the effects of RT-D3 and RT-L4 on Retromer engagement with known regulatory and adaptor proteins.

In contrast to RT-D3, we found that the RT-L4 peptide had little negative impact on the interaction of Retromer with any ligands tested. In GST-pulldowns using purified proteins or using biotinylated peptides and HeLa lysates Retromer was able to interact with TBC1D5, VARP, Fam21, SNX27 and the SNX3-DMT1-II complex in the presence of RT-L4 (**Figs. 4A and 4E**). Interestingly, by ITC we found that the affinity and enthalpy of binding between Retromer for either SNX27_PDZ_, TBC1D5_TBC_, or Fam21_R19-R21_ were substantially improved by the addition of RT-L4 (**Figs. 4B, 4D and 4E; Fig. S7A; Table S5**). Binding between SNX27_PDZ_ and PDZ-dependent cargo peptide, PTHR_586-593_, was also further improved to a *K*_d_ of 700 nM by the presence of RT-L4 (**Fig. 4G**). This suggests that the stabilization of Retromer by RT-L4 may be able to enhance binding to some of its key partners and possibly also improve cargo recognition. However, we did observe a modest reduction in the binding affinity of Retromer for the SNX3-DMT1-II complex from 154 μM to 230 μM in the presence of RT-L4 (**Fig. 4C; Table S6**). We speculate that there may be a small degree of overlap between the binding site for RT-L4 and the first part of the N-terminal loop of SNX3, although not enough to perturb the interaction dramatically. Given this subtle change however, we then asked whether the peptide would perturb the interactions between Retromer and SNX3 in the presence of a lipid membrane. To do this, we performed a liposome binding assay, where we fused a palmitoylated fatty acid to the N-terminus of a cargo peptide. For these experiments we used a sequence derived from the CI-MPR (CIMPR_2347-2376_) and allowed it to insert into liposomes compose of Folch I lipids supplemented with the SNX3-binding phosphatidylinositol-3-phosphate (PtdIns3*P*) (**Fig. 4H; Figs. S7B and S7C**). In our control experiments, we found that SNX3 alone was capable of binding Folch I - PtdIns3*P* liposomes both in the presence and absence of cargo peptide. In contrast, Retromer only bound stably to PtdIns3*P*-containing liposomes when both SNX3 *and* cargo peptides were added, similar to previous studies ^106^. In the presence of RT-L4 we observed a subtle reduction of Retromer binding to SNX3-CI-MPR cargo-PtdIns3*P* liposomes, consistent with the slightly lower affinity observed by ITC (**Fig. S7D**). In summary, peptides binding to Vps29 have a strong and specific impact on a subset of Retromer-associated proteins including TBC1D5 and VARP, without affecting interactions that occur near the Vps35-Vps26 interface (**Fig. 4I**). The stabilising RT-L4 peptide does not prevent binding of any known ligands (**Fig. 4I**), which is an important property of a potential molecular chaperone, although it does have subtle effects on these interactions depending on their respective binding sites.

### RT-L4 reveals an unexpected autoinhibitory role for the C-terminal disordered tails of Vps26A and Vps26B

During our analyses of RT-L4 we noted that it showed a lower binding affinity for the paralogous Vps26B-Vps35_1-390_ subcomplex when compared to Vps26A-Vps35_1-390_ (**Fig. 5A; Table S2**). According to the sequence alignment of Vps26, the residues responsible for Vps35 interaction are highly conserved, while the extended C-terminal tails (residues 298 to 327 in Vps26A) are highly divergent apart from the QRF/YE motif (**Fig 5B; Fig. S7E**). Strikingly when the disordered C-terminal domains of either Vps26A or Vps26B are removed (Vps26A*Δ*C and Vps26B*Δ*C respectively) the binding affinity for RT-L4 is increased to ∼40 nM *K*_d_ and is now identical for both sub-complexes (**Fig. 5A**). We also find a similar improved affinity of RT-L4 for *C. thermophilum* Retromer when the tail of Vps26 is removed (**Fig. S8**). This shows that the C-terminal tails of Vps26A and Vps26B have an autoinhibitory effect on binding to RT-L4, with the Vps26B C-terminus inhibiting the interaction more strongly.

**Figure 5.**
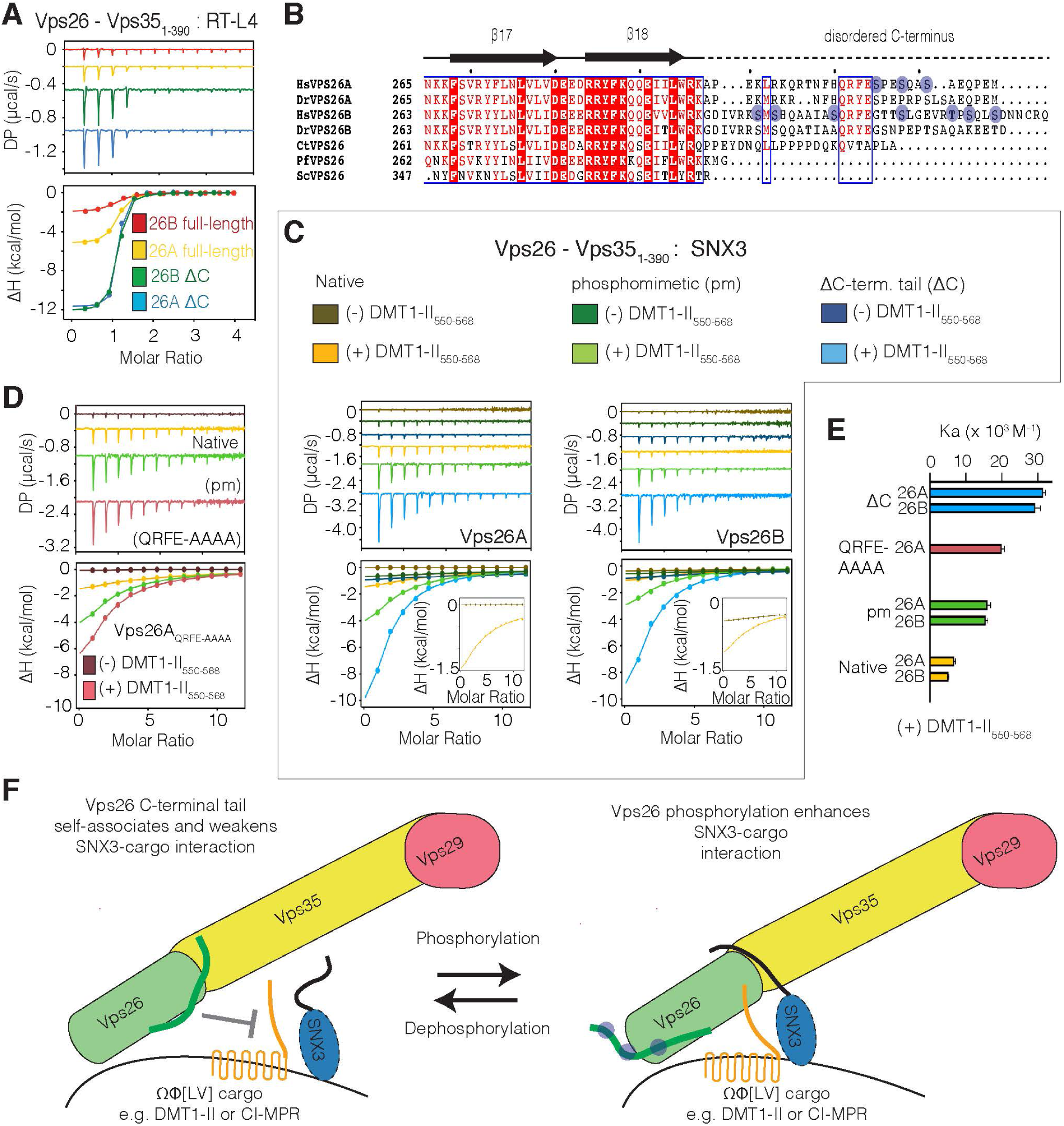
An autoinhibitory role for the disordered C-terminal tail of Vps26 in binding RT-L4 and SNX3. (**A**) ITC thermogram for the titration of RT-L4 with Vps35_1-390_ sub-complex with either full-length or C-terminal truncated Vps26A and Vps26B paralogues. (**B**) Sequence alignment of the C-terminal region of Vps26 highlighting the low-sequence similarity of the unstructured C-terminal tail. Sites of phosphorylation are indicated in blue (www.phosphosite.org) ^107^. Hs, *Homo sapiens*; Dr, *Danio rerio*; Ct, *Chaetomium thermophilum*; Pf, *Plasmodium falciparum*, Sc, *Saccharomyces cerevisiae.* (**C**) ITC measurement of SNX3 binding to native, phosphomimetic (pm), and C-terminal tail truncated (ΔC) versions of Vps26A/B – Vps35_1-390_ subcomplexes. In each case, the presence of DMT1-II cargo peptide was required to detect SNX3 binding. (**D**) ITC measurement of SNX3 binding to QRFE-AAAA mutant Vps26A – Vps35_1-390_. ITC thermograms in (**A**), (**C**) and (**D**) represent the integrated and normalized data fit with a 1 to 1 binding ratio. (**E**) Summary of binding affinities of SNX3 for each Vps26A/B – Vps35_1-390_ subcomplex in the presence of DMT1-II_550-568_ cargo peptide. For clarity, the association constant (*K*_d_^-1^) is shown. The binding affinity is given as mean of at least two independent experiments. (**F**) A proposed model for the autoinhibitory role of the Vps26 disordered C-terminal tails. Our data suggests that these tails can self-associate and reduce affinity for SNX3-cargo complexes, while removal of these tails or their release upon phosphorylation enhances SNX3-cargo association. The C-terminal sequence of Vps26B has greater autoinhibitory activity than Vps26A.

Given that the RT-L4 peptide also subtly affects binding of Retromer to SNX3 and cargos, we were interested to determine whether the C-terminal tails of Vps26 might also influence SNX3 interactions. Intriguingly, SNX3 (in the presence of DMT1-II cargo) binds to Vps26A*Δ*C-Vps35_1-390_ with a significantly increased affinity compared to full-length Vps26A-Vps35_1-390_ (**Fig. 5C; Table S2**). This suggests that there is a self-association of the C-terminal tail that weakens the SNX3-cargo interaction. Previous proteomics have shown that the C-terminal disordered sequences of human and mouse Vps26A and Vps26B can be phosphorylated, although the role of these post-translational modifications (PTMs) and their regulation has not been studied ^107^ (**Fig. 5B)**. We engineered a Vps26A variant with three phosphomimetic mutations in its C-terminal tail and tested the affinity of the Vps35_1-390_/Vps26A phosphomimic complex for SNX3 in the presence of DMT1-II by ITC (**Fig. 5C**). Interestingly this mutant showed an enhanced affinity to the cargo-adaptor complex, although not to quite the extent of the complete removal of the Vps26A tail. Similar enhancement of binding to SNX3 cargo-adaptor complex was also observed when we mutated the conserved QRFE motif of Vps26A to polyalanine (**Figs. 5D and 5E**). Together, our data indicates that the tails of Vps26A and Vps26B play an unexpected autoinhibitory role in the functional interaction of Retromer with the SNX3 adaptor and associated cargos such as Dmt1-II and CI-MPR, and phosphorylation of these tails could activate Retromer to enhance SNX3-cargo interactions (**Fig. 5F**). Vps26B is more strongly autoinhibited than Vps26A, and this observation would explain previous studies showing that the CI-MPR does not bind to Retromer containing Vps26B in cells, but that deletion of the Vps26B tail allows CI-MPR interaction to occur ^108^.

### Macrocyclic peptides as molecular probes to study Retromer mediated endosomal trafficking

With established mechanisms of binding *in vitro* we sought to assess the basic utility of these macrocyclic peptides as novel molecular tools for the study of Retromer. In the first instance we examined their use as fluorescent probes of Retromer localisation in cells (**Fig. 6**). As the peptides were not membrane permeable for cellular uptake we used a reversible cell permeabilization approach with the pore-forming bacterial toxin Streptolysin O (SLO) ^109^. After permeabilization cells were treated with FITC-labelled RT-D3 and RT-L4 and then processed for imaging and co-labelling with specific endosomal markers. In the absence of SLO permeabilization no intracellular fluorescence was detected for the FITC-labelled peptides indicating they are not crossing the membrane or being non-specifically internalised into the lumen of endosomal compartments (**Fig. 6A**). In permeabilised cells however, both FITC-labelled RT-D3 and RT-L4 were recruited to endosomal structures labelled with SNX1 (**Fig. 6A**). Using Airyscan super-resolution microscopy we found that the peptides also co-labelled endosomal structures marked with mCherry-tagged Vps35 (**Fig. 6B**). Lastly, we examined the impact of the peptides on the endosomal recruitment of TBC1D5, which has previously been shown to depend on interaction with Retromer ^71, 76, 77, 95, 110^. As expected, addition of unlabelled RT-L4 had no discernible effect on TBC1D5 localisation; however, RT-L3, which binds to the same site on Vps29 as TBC1D5, caused significant dispersal of TBC1D5 from endosomal structures (**Fig. 6C**). Overall, our data suggests that the Retromer-binding macrocyclic peptides are capable of acting as molecular probes for Retromer localisation.

**Figure 6.**
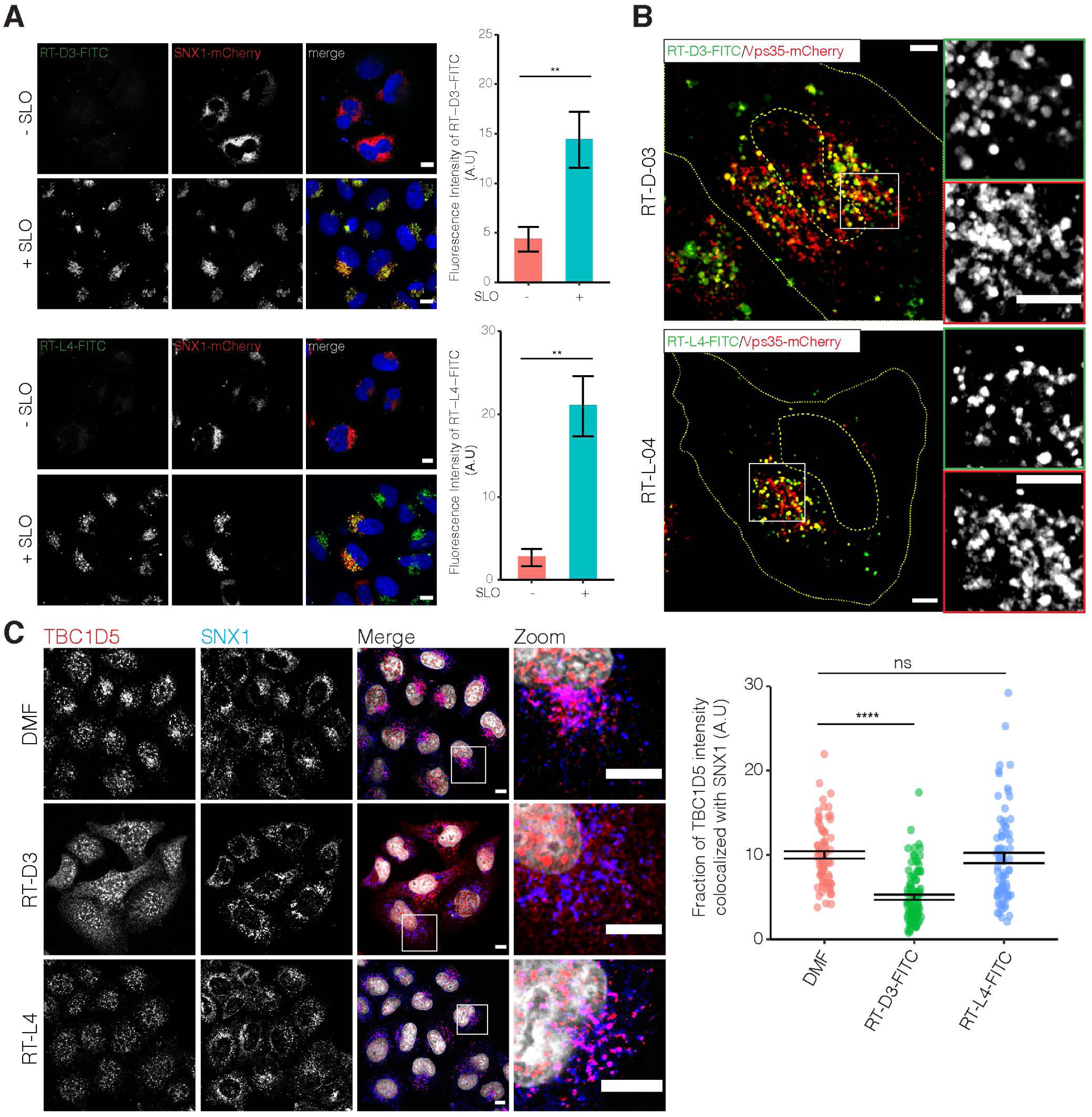
Macrocyclic peptides can be used to study Retromer localization in cells. (**A**) Specific targeting of endosomal structures by streptolysin O (SLO) delivered cyclic peptides. HeLa cells transiently expressing SNX1-mCherry were exposed to SLO at 37°C for 9 min before incubating with the cyclic peptide RT-D3-FITC or RT-L4-FITC on ice for 5 min. Permeabilized cells were recovered in the recovery medium containing Hoechst 33342 for 20 min, then fixed in 4% PFA. The negative control (-SLO) shows no labeling of intracellular structures. Scale bar, 10 μm. Graphs show the fluorescence intensity of RT-D3-FITC or RT-L4-FITC in HeLa cells (means ± SEM). Two-tailed Student’s *t-test* was used to determine the statistical significance (n=3). **, P < 0.01. (**B**) HeLa cells transiently expressing Vps35-mCherry were labeled by SLO-delivered cyclic peptide RT-D3-FITC or RT-L4-FITC, fixed in 4% PFA, and imaged by Airyscan super-resolution microscopy. Scale bar, 5 μm. (**C**) HeLa cells treated with SLO-delivered RT-D3-FITC, RT-L4-FITC, or DMF control were incubated in recovery medium for 2 h, fixed in ice-cold methanol, and co-immunolabeled with antibodies against endogenous TBC1D5 and SNX1, followed by Alexa Fluor-conjugated fluorescent secondary antibodies. Scale bar, 10 μm. The colocalization between TBC1D5 and SNX1 was quantified by Pearson’s correlation coefficient and represented in the graph (means ± SEM). Two-tailed Student’s *t-test* was utilized to determine the statistical significance (n=3). ****, P < 0.0001; ns, not significant.

## DISCUSSION

### De novo macrocyclic peptides that target the Retromer endosomal trafficking complex

In this work the versatile RaPID screen was used to identify a series of potent Retromer-targeting macrocyclic peptides. These bind with high affinity and specificity to Retromer using two distinct evolutionarily conserved binding sites on either Vps29, or at the interface between Vps26 and Vps35. The cyclic peptides in the inhibitor sub-group (RT-D1, RT-D2, RT-D3, RT-L1 and RT-L2) reveal strong binding through the highly conserved surface groove of Vps29, lying on the opposite surface to that bound by the Vps35 subunit. Detailed structural analyses showed that these cyclic peptide inhibitors all form a β-hairpin configuration with a core Pro-Leu motif. Although not tested here, it is likely that these will also be effective at blocking association with the bacterial hijacking molecule RidL, which is known to bind the same site as TBC1D5 and VARP using a similar *β*-hairpin structure ^71, 76, 77^. It is intriguing that all of the Vps29-binding peptides have been selected for the presence of this Pro-Leu dipeptide, and that this peptide has also evolved to mediate binding of endogenous ligands of the Retromer complex such as TBC1D5, VARP and the bacterial effector RidL. The *de novo* macrocyclic peptide screening has therefore inadvertently identified an evolutionarily conserved binding mechanism, and interestingly previous screens of BET domain-binding peptides also uncovered sequence preferences that partly mimicked known endogenous ligands ^111^. Given that Vps29 is also a component of the Retriever complex, a Retromer-related assembly containing homologous subunits Vps35L/C16orf62 and Vps26C/DSCR3 ^112, 113^, these cyclic peptide inhibitors may provide valuable tools for studies of Retriever. Although whether the same binding site in Vps29 is accessible within the Retriever complex remains to be confirmed.

Among the identified macrocyclic peptides RT-L4 binds specifically to the interface between the N-terminal domain of Vps35 and the C-terminal sub-domain of Vps26 and is the sole molecule to show a significant ability to stabilise the Retromer complex. In the recent cryoEM structure of mammalian Retromer it was found that Vps26 and the N-terminal portion of Vps35 exhibit substantial flexibility compared with other regions of the complex ^100^. We speculate that the improved thermal stability of Retromer in the presence of RT-L4 may be partly due to reduced flexibility in these two subunits. Importantly, the RT-L4 peptide did not inhibit the binding of Retromer to its essential interacting partners including SNX27, Fam21, TBC1D5 and VARP, and in fact led to a general increase in their binding affinity *in vitro*. RT-L4 did have a minor effect however on binding of Retromer to SNX3 and *ΩΦ*[LV] motif-containing cargo, consistent with the fact that the RT-L4 binding site partially overlaps with the site where the N-terminal disordered domain of SNX3 engages Vps26 and Vps35 ^98^.

### Macrocyclic peptides as molecular tools for the study of Retromer function

The series of macrocyclic peptides discovered here provide novel molecular probes for the study of Retromer and endosomal trafficking. Firstly, using a simple reversible permeabilization approach, we have successfully delivered the cyclic peptides RT-D3 and RT-L4 into cells and find they are specifically targeted to the Retromer-positive endosomal structures. Thus, the peptides can be used to examine endogenous Retromer localisation *in situ* and could feasibly be used to study Retromer in live cells or using super-resolution approaches with different types of fluorescent dyes apart from the simple FITC labelling strategy used here. The RT-D3 peptide also impacts recruitment of proteins such as TBC1D5 that depend on binding to the Vps29 subunit of the complex, and therefore could be used to probe the impact of acutely disrupting this interaction in the future. In addition to studying the localisation of Retromer, the RT-L4 peptide provided a highly specific substrate for purification of the Retromer complex from cells, and in the future could be useful for proteomic studies of Retromer in diverse cells, tissues, and organisms. We also envisage that these peptides will provide tools for enhancing Retromer stability for future structural studies of its interactions with accessory and regulatory proteins by cryoEM and X-ray crystallography.

Our studies of the RT-L4 peptide binding to the Vps26–Vps35 interface revealed an autoinhibitory role for the C-terminus of the Vps26A and Vps26B paralogues. Removal of their divergent C-terminal tails significantly increased the affinity of both Retromer complexes for the RT-L4 peptide to similar levels, leading us to then test the impact of the Vps26A and Vps26B tail domains on binding to the SNX3–DMT1-II cargo-adaptor complex, a native ligand that engages Retromer at a similar location. Again, removal of the C-terminal disordered domains significantly improved the affinity of Retromer for SNX3–DMT1-II, confirming their autoinhibitory activity has a functional importance. Furthermore, this may be regulated by phosphorylation of the Vps26A and Vps26B tails, as phosphomimetic mutants have a higher affinity for SNX3-DMT1-II than the wild-type proteins. Since the discovery of the Vps26B paralogue of Vps26A it was clear that the most significant difference between the two proteins was the sequence of their C-terminal domains ^4, 114^. Later it was shown that Vps26A but not Vps26B Retromer bound to the SNX3-dependent cargo CI-MPR, however, CI-MPR binding and trafficking by Vps26B was restored when its tail was deleted ^108^. Our data demonstrates a direct role for both the Vps26A and Vps26B tails in negatively regulating the binding of SNX3-cargo *in vitro*, with the Vps26B tail possessing a more potent autoinhibitory sequence than that of Vps26A. This suggests that an activating switch such as phosphorylation of Vps26 proteins or binding of another regulatory protein may promote the recruitment of SNX3-cargo complexes by Retromer in cells, although the specific nature of this switch remains to be determined.

### Towards therapeutic targeting of Retromer

Endolysosomal trafficking and regulation of proteostasis is emerging as an attractive target in the treatment of a range of diseases, including neurodegenerative disorders like AD and PD, infection by viral and bacterial pathogens, and other diseases impacted by defective endosomal signalling such as cancer. There is an increasing interest in developing molecules for both inhibition and enhancement of Retromer activity in these processes ^20, 39, 40^. As shown previously, peptides that target Retromer and compete with the viral L2 protein can reduce HPV infection in cell and animal models ^66, 78, 79^, while a small molecule with Retromer chaperoning activity can reduce cellular accumulation of toxic material causing neurodegeneration ^54, 80–90^. However, these peptides and small molecules have only a relatively low affinity for Retromer, and their specificities and pharmacological profiles are essentially unknown.

Because of their typically high affinity and larger surface area coverage, macrocyclic peptides are emerging as an important class of molecules for the design of new drugs and molecular probes targeting proteins and protein-protein interactions, often considered difficult using traditional small molecule approaches ^93, 115–118^. The peptides we have discovered can be classified as either Retromer inhibitors or stabilizers, and we provide a comprehensive biochemical and structural explanation for how each cyclic peptide associates with Retromer and affects its native molecular interactions. The Retromer inhibitors we identified were able to potently block TBC1D5 and VARP binding and will likely also preclude interaction of the bacterial effector RidL, suggesting a conserved site in Retromer primed for targeting by peptide or small molecule-based inhibition. The stabilising peptide RT-L4 enhanced Retromer association with known regulatory proteins, and its mechanism of action and superior affinity to the R55 small molecule highlights the potential of targeting the Vps26 and Vps35 interface for the design of novel pharmacological chaperones of Retromer in future studies. While the macrocyclic peptides described here are potential leads to therapeutic targeting of Retromer, notable hurdles include cell permeability, oral availability, and an ability to cross the blood brain barrier when targeting neurological diseases. However, new approaches show promise in overcoming these barriers, including the use of non-standard amino acids with novel activities and solubility profiles, coupling to various cell-targeting peptides, and novel delivery methods ^118–121^. Alternatively, the peptides discovered here could provide the basis for competitive screening for drug-like molecules that target the same binding sites.

In summary, we have identified a series of Retromer-targeting macrocyclic peptides and demonstrate their potential for Retromer inhibition and activation based on a comprehensive understanding of their different mechanisms of action. They will be a valuable resource for the study of Retromer function at the cellular and molecular level and represent promising leads for the targeting of Retromer in a variety of diseases caused by dysregulation or disruption of the endosomal membrane trafficking system.

## MATERIALS AND METHODS

### Chemicals and antibodies

Rabbit polyclonal anti-TBC1D5 was purchased from Proteintech (17078-1-AP). Mouse monoclonal anti-SNX1 was purchased from BD Transduction Laboratories (611483). Mouse monoclonal anti–α-tubulin (clone DM1A; T9026) was purchased from Sigma-Aldrich. Goat polyclonal anti-Vps35 (NB100-1397) was purchased from Novus Biologicals. Secondary donkey anti-mouse IgG Alexa Fluor 647 (A31571), and donkey anti-rabbit IgG Alexa Fluor 555 (A31572) were purchased from Thermo Fisher Scientific. The linear peptides DMT1-II_550-568_ (AQPELYLLNTMDADSLVSR) and palmitoylated CI-MPR_2347-2375_ with N-terminal di-lysine (KKSNVSYKYSKVNKEEETDENETEWLMEEIQ) were both synthesised by Genscript (USA).

### Molecular Biology and Cloning

pmCherry-SNX1 was generated by subcloning the full-length open reading frame of human SNX1 from pEGFP-SNX1 described previously ^122^, into the multiple cloning site of pmCherry-C1. pmCherry-Vps35 was generated by subcloning the full-length open reading frame of human Vps35 from pEGFP-Vps35 described previously ^18^, into the multiple cloning site of pmCherry-N1.

For bacterial expression, Retromer constructs encoding full-length human and zebrafish Vps29, Vps26A and Vps35 were cloned into either pET28a and pGEX4T-2 vectors as described previously ^99, 123^. *Chaetomium thermophilum* Vps29, Vps26 and Vps35 were also cloned using protocol described previously ^97^. In all cases, Vps26A was cloned as a N-terminal His_6_-tag fusion protein and Vps29 was cloned into pGEX4T-2 vector as a cleavable N-terminal GST fusion protein. For mouse Vps26B, full-length cDNA was inserted into pMW172Kan vector ^114^. For the truncation constructs, the DNA sequence encoding the N-terminal part of Vps35 (Vps35_1-172_, Vps35_1-224_ and Vps35_1-390_) was cloned into pGEX4T-2 vector. Vps26 ΔC-term. tail (ΔC) constructs (Vps26A_9-298_ and Vps26B_7-296_) were cloned into pET28a vector containing His_6_-tag. Full-length Vps26A pm mutant (substituted S315E, S318E and S321E), Vps26A_QRFE-AAAA_ (substituted residues 311 - 314 to alanine) and Vps26B pm mutant (S302E, S304E, S311E, S319E, T325E, S327E, and S330E substitution) were synthetic genes by Genscript Corporation. cDNA encoding full-length human SNX3 was cloned into pGEX4T-2 vector. Similarly, the cDNA encoding the human Fam21 LFa motif repeats 19 to 21 was cloned into pGEX4T-2 vector. The TBC domain of human TBC1D5 was cloned into the pMCSG9 vector containing a N-terminal GST and a TEV cleavage site. Full-length mouse SNX27 was cloned into the pMCSG7 vector containing a N-terminal His and a TEV cleavage site ^124^. PDZ domain of mouse SNX27 was cloned into the pGEX4T2 vector similar to the one described previously ^14^. Site-directed mutagenesis was performed to generate mutant constructs with custom-designed primers. All constructs were verified using DNA sequencing.

### RaPID screening

For the first round of RAPID selection, an mRNA library was generated by T7 polymerase mediated *in vitro* transcription of a PCR assembled DNA template, purified by PAGE, and covalently ligated to a puromycin linked oligonucleotide with T4 RNA ligase. 1.2 μM puro-linked mRNA library was translated in a 150 μL *in vitro* translation reaction (genetically reprogrammed to incorporate L- or D- ClAC-Tyr in place of the initiator formyl-methionine) at 37°C for 30 min. Peptide-mRNA fusion molecules were released from the ribosome by treatment with 17 mM EDTA for 30 min. at 37°C, reverse transcribed using MMLV reverse transcriptase (H-)(Promega) at 42°C for 1 hour and buffer exchanged to TBS-T using sephadex G-25. The resulting cyclic peptide-mRNA:cDNA library was counter-selected 3 times against His Pull-Down Dynabeads (Thermo Fisher), and affinity selected against 200 nM bead-immobilised Retromer for 30 min. at 4°C, with the beads washed 3 times with TBS-T, overlaid with 0.1% triton-X100 and heated to 95°C to elute the cDNA for recovery by PCR.

For the second and subsequent rounds of selection, the translation reaction was scaled down to 2.5 μL total volume, and 6 iterative counter-selections using uncoated beads were conducted prior to affinity selection against Retromer. Following 5 iterative rounds of selection, Retromer ligands were identified by sequencing the final enriched cDNA using a MiSeq sequencer (Illumina).

### Synthesis of Cyclic Peptides

Untagged peptides were synthesized at 25 μM scale on NovaPEG Rink Amide resin (0.53 mmol/g) using Fmoc-based chemistry on a Syro I peptide synthesizer (Biotage). Fmoc-protected amino acid (6 eq.), (2-(1H-benzotriazol-1-yl)-1,1,3,3-tetramethyluronium hexafluorophosphate (HBTU, 6 eq.), hydroxybenzotriazole (HOBt, 6 eq.) and disopropylethylamine (DIPEA, 12 eq.) with 30 min coupling cycles. Deprotection of fmoc was achieved with 40% piperidine/DMF for 1 × 3 min. then 1 x 12 min. Chloroacetic acid was coupled manually after the final deprotection using the same conditions.

Linear peptides were cleaved from resin with TFA/TIS/H_2_O/EDT (92.5:2.5:2.5:2.5) over 2 h. Crude peptides were precipitated and washed (5x cold Et_2_O), redissolved in DMSO, and cyclized by adding DIPEA until basic followed by incubating at room temperature. Cyclic peptides were acidified, diluted to 50% DMSO with water and purified by RP-HPLC using a Chromolith C18 column with a gradient of 10 to 70% buffer B (99.9% CH_3_CN/0.1% TFA in buffer A, 0.1% TFA in water) over 40 min and lyophilized, before the TFA salt was exchanged to HCl by triplicate lyophilization from 5 mM HCl aq.

### Synthesis of biotinylated and FITC labelled cyclic peptides

Biotin and FITC labelled peptides were synthesized (100 μM scale) on Rink Amide MBHA resin (0.6 mmol/g) using Fmoc-based chemistry and a peptide synthesizer (Symphony, Protein Technologies) Fmoc -Lys(Mtt)- OH in position 1. Fmoc-protected amino acid (4 eq.), 2-(6-chloro-1H-benzotriazole-1-yl)-1,1,3,3 tetramethylaminium hexafluorophosphate (HCTU, 4 eq.), and disopropylethylamine (DIPEA, 4 eq.) were used in 2 × 30 min coupling cycles. Fmoc deprotection was achieved by treatment with 1:2 piperidine/DMF for 2 × 3 min. Chloroacetic acid was coupled manually after the final deprotection using chloroacetic acid (4 eq.) and HATU (4 eq.), and DIPEA (4 eq.) in N,N-dimethylformamide (DMF). Selective side-chain deprotection of the methyl trityl (Mtt) group on lysine was achieved using 3% TFA in dichloromethane (DCM) (5 *×* 2 min). Biotin was coupled using biotin 2-(1H-7-azabenzotriazol-1-yl)-1,1,3,3-tetramethyl uraniumhexafluorophosphate methanaminium (HATU, 2 eq.) and DIPEA in DMF. FITC was coupled using FITC (2 eq.) and DIPEA.

Linear peptides were cleaved from the resin by treatment with TFA/TIS/H_2_O (95:2.5:2.5) for 2 h. The crude peptides were precipitated and washed with cold Et_2_O, redissolved in 50% acetonitrile/0.05% TFA in water, and lyophilized. Peptides were purified by RP-HPLC using a Phenomenex Luna C18 column eluting at a flow rate of 20 mL/min and a gradient of 20 to 70% buffer B (90% CH_3_CN/10% H_2_O/0.1% TFA in buffer A, 0.1% TFA in water) over 30 min and lyophilized.

Cyclization to form the thioether was achieved by dissolving the linear peptide in DMF with DIPEA (10 eq.) and the reaction monitored by UP-LCMS. After no more linear peptide was detected, DMF was removed *in vacuo* and the cyclic peptide purified by RP-HPLC using a Phenomenex Luna C18 column eluting at a flow rate of 20 mL/min and a gradient of 20 to 70% buffer B (90% CH_3_CN/10% H_2_O/0.1% TFA in buffer A, 0.1% TFA in water) over 30 min and lyophilized.

Peptides were characterised and purity determined by analytical RP-UPLC and UPLC-MS methods. UPLC was performed on Shimadzu Nexre UPLC with PDA using an AgilentEclipse plus C18RRHD 1.8 μm, 2.1 ′100 mm UPLC Column., and a gradient of 0 to 80% buffer B (90% CH_3_CN/10% H_2_O/0.1% TFA in buffer A, 0.1% TFA in water) over 6 min. UPLC-MS was performed on Shimadzu Nexre UPLC system connected to LCMS-2020 single quadrupole mass spectrometer using an phenomonex Aeris Peptide 1.7 μm XB-C18 column 50 x 2.1 mm and a gradient of 0 to 80% buffer B (90% CH_3_CN/10% H_2_O/0.1% formic acid in buffer A, 0.1% formic acid in water) over 6 min.

### Cell culture and transfection

HeLa cells (ATCC CCL-2) were maintained in high glucose Dulbecco’s Modified Eagle Medium (DMEM; Thermo Fisher Scientific) supplemented with 10% fetal bovine serum (FBS) and 2 mM L-glutamine (Thermo Fisher Scientific) in a humidified 37 °C incubator with 5% CO_2_. Transfection was performed using Lipofectamine 2000 (Thermo Fisher Scientific) according to the manufacturer’s instructions.

### Reversible permeabilization and peptide delivery

Reversible permeabilization of HeLa cells with Streptolysin O (SLO; 25,000 - 50,000 U; Sigma-Aldrich) was performed as previously described ^125^. In brief, an aliquot of SLO stock was reduced by 10 mM DTT (Sigma Aldrich) for 20 min at 37°C, then diluted to working concentration (200x) in DPBS containing 1 mM MgCl_2_. HeLa cells grown on coverslips were incubated in the SLO containing solution for 9 min at 37°C, washed twice with DPBS containing 1 mM MgCl_2_, then incubated with ice-cold transport buffer (140 mM NaCl, 5 mM KCl, 1 mM MgCl_2_, 10 mM HEPES, 10 mM glucose, pH 7.4) containing fluorescent peptides (5 µg/ml) on ice for 5 min. After labeling, the cells were washed with ice-cold transport buffer twice and incubated with the recovery medium (DMEM, 10% FBS, 1.8 mM CaCl_2_, without antibiotics) for 20 min at 37 °C. Cells were fixed with 4% PFA or ice-cold methanol and subjected for microscopy analysis.

### Indirect immunofluorescence and image analysis

HeLa cells grown on coverslips were routinely fixed and permeabilized in ice-cold methanol for 5 min at −20°C, unless otherwise cited. After blocked with 2% BSA in PBS for 30 min, cells were labeled with anti-TBC1D5 (1:200) and anti-SNX1 (1:100) for 1 hr at room temperature followed by the incubation with Alexa Fluor 555 and 647 conjugated secondary antibodies. Coverslips were mounted on glass microscope slides using Fluorescent Mounting Medium (Dako), and the images were taken at room temperature using the Leica DMi8 SP8 inverted confocal equipped with 63x Plan Apochromatic objectives or the Zeiss LSM880 Axiovert 200 inverted confocal with AiryScan FAST detector equipped with 63x Plan Apochromatic objectives. For quantification, images were taken from multiple random positions for each sample.

Images were processed using ImageJ software. Colocalization analysis was performed as described previously ^12^. In brief, single cells were segregated from fields of view by generating regions of interest, cropped, split into separated channels, and applied for threshold processing. Colocalization analysis was conducted on three independent experiments and represented as Pearson’s correlation coefficient. Colocalization values were exported to R studio and tabulated accordingly.

### Recombinant Protein Expression and Purification

All the Retromer constructs were expressed using BL21 Star^TM^ (DE3) cells and induced by the addition of IPTG to a final concentration of 1 mM at an OD_600_ of ∼ 0.8. The temperature was then reduced to 18°C and incubated overnight. To obtain the Retromer trimer complex (including human, zebrafish and *Chaetomium thermophilum*), full-length GST-tagged Vps29 co-expressed with Vps35 was mixed with cell pellet of Vps26A and lysed through a Constant System TS-Series cell disruptor in lysis buffer containing 50 mM HEPES pH 7.5, 200 mM NaCl, 5% glycerol, 50 μg/ml benzamidine and DNase I. The homogenate was cleared by centrifugation and loaded onto Talon® resin (Clonetech) for purification. To obtain the correct stoichiometry ratio of the Retromer complex, the purified elution from Talon® resin was further passed through the glutathione sepharose (GE healthcare). Removal of the GST-tag from human and zebrafish Vps29 was performed using on-column cleavage with thrombin (Sigma-Aldrich) overnight at 4°C. For *Chaetomium thermophilum* Vps29, the GST-tag was cleaved using PreScission protease overnight at 4°C. The flow-through containing Retromer complex with GST-tag removed was further purified using size-exclusion chromatography (SEC) on a Superdex200 column equilibrated with a buffer containing 50 mM HEPES pH 7.5, 200 mM NaCl, 5% glycerol and 0.5 mM TCEP. Production of Retromer individual subunits (Vps29, Vps26A and Vps35) and subcomplexes (Vps29 - Vps35, Vps26A - Vps35_1-172_, Vps26A - Vps35_1-224_, Vps26A - Vps35_1-390,_ Vps26A_ΔC-term. tail_ - Vps35_1-390_, Vps26A_pm_ - Vps35_1-390_, Vps26A_QRFE-AAAA_ - Vps35_1-390_, Vps26B - Vps35_1-390_, Vps26B_ΔC-term. tail_ - Vps35_1-390_, and Vps26B_pm_ - Vps35_1-390_) were expressed and purified under the same method. In brief, glutathione sepharose was used for Vps29, Vps35 and subcomplexes purification, and Talon® resin was applied for Vps26A.

Expression and purification of GST-tagged SNX3, GST-tagged Fam21_R19-R21_, TBC1D5_TBC_, SNX27_PDZ_ and His-tagged SNX27 were performed using similar methods described previously ^14, 95, 98, 124^. Cell pellets were lysed by Constant System TS-Series cell disruptor using the same buffer as Retromer. All proteins were passed through either Talon® resin or Glutathione sepharose depending on the affinity tag. Removal of fusion tag was performed on-column overnight using either thrombin for SNX3, and TEV protease for SNX27_PDZ_ and TBC1D5_TBC_. The flow-through containing GST-tag removed SNX3, TBC1D5 and SNX27_PDZ_ were further purified using SEC in the same way as Retromer. For, His-tagged SNX27, the fractions eluted from Talon® resin was further purified SEC directly using the same buffer as described above. Similarly, the GST-tagged Fam21_R19-R21_ eluted from Glutathione sepharose was directly injected into SEC for final purification.

Retromer 3KE construct design, expression, and purification has previously been described ^100^. Briefly, Retromer plasmids were co-transformed into BL21(DE3) Rosetta2 pLysS cells (Millipore). Cells were grown to OD_600_ between 0.8-1.0 and induced for 16-20 hours at 20°C with 0.4 mM IPTG. Cells were lysed by a disruptor (Constant Systems Limited). Protein was purified in 10 mM Tris-HCl (pH 8.0), 200 mM NaCl, 2 mM *β*ME using glutathione sepharose (GE Healthcare). Protein was cleaved overnight using thrombin (Recothrom, The Medicines Company) at room temperature and batch eluted in buffer. Retromer was further purified by gel filtration on a Superdex S200 10/300 column (GE Healthcare) into 10 mM Tris-HCl (pH 8.0), 200 mM NaCl.

### Biotinylated Cyclic Peptides Pull-down Assay

Culture dishes (15 cm) with HeLa cells at approximately 90% confluency were washed with PBS and lyzed by the lysis buffer containing 50 mM HEPES pH 7.5, 200 mM NaCl, 1% Triton, 25 µg/ml DNase I and one protease-inhibitor cocktail tablet per 50 ml lysis buffer. Soluble and insoluble fractions were separated by centrifugation at 13,000 *g* for 20 min at 4°C. After centrifugation, supernatant was then added to the streptavidin agarose (Thermo Scientific) pre-incubated with 140 µM of either biotinylated RT-D3 or biotinylated RT-L4 cyclic peptides for 2 h at 4°C. Both cyclic peptides were carefully prepared without forming precipitation before mixing with streptavidin agarose.

In the case of capturing SNX3, roughly 5 µM of DMT1-II_550-568_ peptide was added to the supernatant prior mixing with the streptavidin agarose. Beads were then spun down at 2,000 g for 2 min and washed five times with washing buffer containing 50 mM HEPES pH 7.5, 200 mM NaCl, 5% glycerol, 0.05% triton X-100 and 0.5 mM TCEP. Bound complex was eluted from the streptavidin agarose by boiling in 100 mM DTT added SDS loading buffer (Life Science) and subjected to SDS-PAGE analysis and western blotting.

### RT-L4 Binding Site Screening Assay

Mapping the potential binding region of RT-L4 was carried out using purified Vps26A, Vps35_1-390_, Vps26-Vps35_1-390_ subcomplex and the associated mutants. First, 10 µM of purified proteins were incubated with fresh streptavidin agarose containing either 100 µM of RT-L4 or equivalent percentage (v/v) of DMSO. The mixture was incubated in binding buffer containing 50 mM HEPES pH 7.5, 200 mM NaCl, 5% glycerol, 0.5 mM TCEP for 30 min at 4°C. Beads were then washed three times with binding buffers followed by SDS-PAGE analysis.

### GST Pull-down Assay

GST pull-down assay was carried out using either GST-tagged Retromer, GST-Fam21_R19-R21_, or GST- TBC1D5_TBC_ as bait protein. For pull-down assays containing either SNX27 or SNX3 + DMT1-II_550-568_ peptide, GST-tagged Retromer and GST alone were first incubated with fresh glutathione sepharose bead for 2 hours at 4°C. To avoid precipitation caused by cyclic peptides, SNX3/SNX27 - cyclic peptides mixture were centrifuged at 17,000 rpm for 20 min at 4°C before added into the glutathione sepharose bead samples. The reaction mixtures were incubated for at least 4 hours in binding buffer containing 50 mM HEPES pH 7.5, 200 mM NaCl, 5% glycerol, 0.1% triton X-100 and 0.5 mM TCEP. Beads were then washed four times with binding buffers and samples of beads were analyzed by SDS-PAGE. Retromer - TBC1D5 and Retromer – Fam21 pull-down assay were performed using identical protocol as described above. For Retromer - TBC1D5 pull-down assay, GST-TBC1D5_TBC_ was used in order to differentiate GST-Vps29 and TBC1D5_TBC_ on the SDS-PAGE.

### Crystallization and Data Collection

Crystallization screening was performed using hanging drop vapour diffusion method under 96-well format at 20°C. To co-crystallize Vps29 with cyclic peptides, a 1.5-fold molar excess of the RT-D1, RT-D2, RT-D3, RT-L1 and RT-L2 peptides were added separately to the purified hVps29 to a final concentration of 14 mg/ml. Initial crystals were obtained in hVps29 - RT-D1, hVps29 - RT-D2 and hVps29 - RT-L2 complex samples. For hVps29 - RT-D1, plate shape crystals were observed in many different commercial screen conditions, but the best quality crystals were obtained in a condition comprising 3.5 M sodium formate. For hVps29 – RT- D2, the optimized crystals were obtained by streak-seeding crystals grown in 0.1 M potassium thiocyanate, 30% PEG 2000 MME into the same condition prepared with protein at 8 mg/ml. For hVps29 – RT-L2 sample, precipitation was observed after the addition of the cyclic peptide. Precipitation was removed by centrifugation and diamond shape crystals were obtained after overnight incubation in condition consisted of 0.1 M HEPES pH 7.0, 1 M succinic and 1% PEG2000 MME. Initial attempts to co-crystallize hVps29 – RT-D3 and hVps29 – RT-L1 were unsuccessful. For hVps29 – RT-L1, small long needle shape crystals were observed in condition consisted of 1.4 M sodium malonate pH 6.0 using 26 mg/ml protein with 1.5 fold molar excess of RT-L1 and 10 fold molar excess of 18-crown-6. Diffraction quality crystals were obtained by streak-seeding crystals grown in the 26 mg/ml into the same condition prepared with protein at 15.5 mg/ml. To grow crystals containing RT-D3, we substituted hVps29 to ctVps29, and managed to obtain diffraction quality crystals in condition consisted of 0.18 M ammonium citrate dibasic and 20% PEG 3350 using 14 mg/ml protein with 2- fold molar excess RT-D3. Prior to data collection, all the crystals were soaked in the appropriate cryoprotectant solutions. X-ray diffraction data were measured on the MX1 and MX2 beamlines at the Australian Synchrotron at 100 K.

### Crystal Structure Determination

All the data collected were indexed and integrated by AutoXDS ^126^ and scaled using Aimless ^127^. Crystal structures of hVps29 - RT-D1, hVps29 - RT-D2, ctVps29 - RT-D3, hVps29 - RT-L1 and hVps29 - RT-L2 were solved by molecular replacement using Phaser ^128^ with native hVps29 crystal structure (PDB ID: 1W24) as the initial model. The initial electron density map obtained from the best solution guided the locations of the cyclic peptides and 18-crown-6. Structure refinement was performed using the PHENIX suite ^129^ with iterative rebuilding of the model. The refined model was manually rebuilded using Coot guided by *F_o_ - F_c_* difference maps. Coordinates for D-tyrosine and sulfanylacetic acid linking N-terminal and C-terminal of the peptides were generated using LIBCHECK from Coot. The quality and geometry of the refined structures were evaluated using MolProbity ^130^. Data collection and refinement statistics are summarized in **Table S3**. Sequence conservation was based on a *T-Coffee* multiple sequence alignment ^131^ and calculated using the ConSurf Server ^132^. Structure comparison was analysed using DALI ^133^ and molecular figures were generated using PyMOL.

### Surface Plasmon Resonance

SPR experiments were conducted at room temperature using a Biacore T-200 instrument (Cytiva) in HBS-P+ buffer (10 mM HEPES, 150 mM NaCl, 0.05 % (v/v) surfactant P20, pH 7.4) containing 0.1% (v/v) DMSO. 100 nM human or zebrafish Retromer was immobilized on a Ni^2+^-primed Series S Sensor Chip NTA (Cytiva) following the manufacturer’s instructions. A single cycle kinetics protocol involving five 120 s injections of peptide as analyte at a flow rate 60 μL.min^-1^ was employed, with kinetics determined using a 1:1 binding model.

### Isothermal Titration Calorimetry

ITC experiments were conducted at 25°C using a Microcal ITC200 (Malvern) in buffer containing 50 mM HEPES pH 7.4, 200 mM NaCl, 5% glycerol, 0.5 mM TCEP. Cyclic peptides in the range of 120 μM to 300 μM were titrated into 6 – 20 μM of Retromer, subcomplexes or individual subunits. The interaction of Retromer and TBC1D5_TBC_ in the presence of cyclic peptide was carried out by titrating 80 μM of TBC1D5_TBC_ into 6 μM of Retromer + 30 μM of either RT-D3 or RT-L4. In the native control, the cyclic peptide was substituted with equivalent percentage (v/v) of DMSO. Similarly, the effect of the cyclic peptides on the interaction of Retromer and SNX27_PDZ_ was performed by titrating 1.3 mM of SNX27_PDZ_ into 30 μM of Retromer + 150 μM of either RT-D3 or RT-L4. The interaction of Retromer and Fam21 was performed by titrating 300 μM of GST-Fam21_R19-R21_ into 12 μM of Retromer + 60 μM of either RT-D3 or RT-L4. In the case of SNX3, 1.2 mM SNX3 was titrated into 12 μM of Retromer + 180 μM DMT1-II_550-568_ peptide + 60 μM of either RT-D3 or RT-L4. Similarly, 1.2 mM SNX3 was titrated into 12 μM of Retromer + 180 μM CIMPR_2347-2376_ peptide to examine its effect on Retromer and SNX3 interaction. To ensure that RT-L4 binds specifically only to Retromer but not the accessory proteins, 90 μM of RT-L4 was titrated into 7 μM Retromer, TBC1D5_TBC_, SNX3 or SNX27 using buffer described above. The effect of Vps26 C-terminal disordered tail on RT-L4 and SNX3 binding was first performed by titrating 90 μM of RT-L4 into 8 μM of Vps26A/B - Vps35_1-390_, or Vps26A/B_ΔC-term. tail_ - Vps35_1-390_ subcomplexes. In the case of SNX3 binding, 970 μM of SNX3 was titrated into 17 μM of FL, pm mutant, QRFE-AAAA or ΔC Vps26A/B - Vps35_1-390_ subcomplexes with and without 255 μM DMT1-II_550-568_ peptide. For the Retromer and R55 interaction, 120 μM - 640 μM of R55 was titrated into 9 μM – 16 μM of Retromer using the same buffer as the cyclic peptide ITC experiments.

In all cases, the experiments were performed with an initial 0.4 μl (not used in data processing) followed by 12 serial injections of 3.22 μl each with 180 sec intervals. Data were analyzed with Malvern software package by fitting and normalized data to a single-site binding model, yielding the thermodynamic parameters *K*_d_, Δ*H*, Δ*G* and -TΔ*S* for all binding experiments. The stoichiometry was refined initially, and if the value was close to 1, then N was set to exactly 1.0 for calculation. All experiments were performed at least in triplicate to check for reproducibility of the data.

### Differential Scanning Fluorimetry

Thermal unfolding experiments were carried out through preferential binding of a fluorophore to unfolded protein using a ViiA7 real-time PCR instrument (Applied Biosystems). In brief, 0.4 mg/ml of fresh Retromer, subcomplex and individual subunits was pre-incubated with 30 - 60 molar excess of cyclic peptides for at least 30 min on ice followed by centrifugation at 17,000 rpm for 20 min at 4°C to remove all possible precipitation. To measure thermal denaturation, freshly prepared SYPRO orange dye (Life Science) was then added to protein-cyclic peptide complex mixture to a final concentration of 5X before loaded into the 96-well plate. Relative fluorescence units (R.F.U.) were measured from 25°C to 90°C using the ROX dye calibration setting at 1°C increments. Experiments were performed with four replicates and T_m_ was calculated using Boltzmann sigmoidal in Prism version 8.0.1 (GraphPad software).

### Mass Photometry

Molecular mass measurement of Retromer in the presence of cyclic peptide was performed using Refeyn OneMP mass photometer (Refeyn Ltd). In brief, 10 µl of standard buffer containing 50 mM HEPES pH 7.5, 200 mM NaCl was applied. Next, 1 µl of 50 nM Retromer + RT-D3 & RT-L4 cyclic peptide was added to the drop to a final concentration of 5 nM and 10000 frames were recorded. Calibration was performed using three protein standards (i.e. 66, 146 and 480 kDa) (ThermoFisher Scientific).

### Cryo-EM Grid Preparation and Data Collection

For cryo-electron microscopy of Retromer+RT-L4, Retromer 3KE at a final concentration of 0.5 mM in 20 mM Tris pH 8.0 / 100 mM NaCl / 2 mM DTT was combined with RT-L4 at a final concentration of 0.1 mM, incubated for 1 hour, and spun briefly in a tabletop centrifuge. 2 μl of the sample was applied to freshly glow discharged Quantifoil 1.2/1.3 300 mesh grids, and the grids were vitrified in liquid ethane using a ThermoFisher Mark IV Vitrobot, using a 3.5 second blot time at 100% humidity and 20°C. 4791 micrographs were collected on a ThermoFisher FEI Titan Krios G3i microscope in the Center for Structural Biology’s Cryo-EM Facility at Vanderbilt. The microscope operated at 300 keV and was equipped with a ThermoFisher Falcon3 direct electron detector camera. The nominal magnification used during data collection was 120,000x, and the pixel size was 0.6811 Å/pix. The total electron dose was 50 e^-^/A^2^, and micrographs were collected at +/-30° tilts. Data collection was accomplished using EPU (ThermoFisher).

For cryo-electron microscopy of *apo* Retromer, 2 μl WT Retromer at a concentration of 0.5 mM in 20 mM Tris pH 8.0 / 100 mM NaCl / 2 mM DTT was applied to freshly glow discharged Quantifoil 1.2/1.3 300 mesh grids, and the grids were vitrified in liquid ethane using a ThermoFisher Mark IV Vitrobot, using a 2s blot time at 100% humidity and 8°C. 891 micrographs were collected on a ThermoFisher FEI Titan Krios microscope at the National Resource for Automated Molecular Microscopy (NRAMM). The microscope operated at 300 keV and was equipped with a Gatan BioQuantum energy filter with a slit width of 20eV and a Gatan K2 Summit direct electron detector camera. The nominal magnification used during data collection was 105,000x, and the pixel size was 1.0691 Å/pix. The total electron dose was 73.92e^-^/A^2^, and micrographs were collected at +/-15° tilts. Data collection was accomplished using Leginon ^134^

### Single Particle Cryo-EM Image & Data Processing

All images were motion corrected using MotionCor2 (Zheng et al., 2017). Micrographs from the *apo* Retromer data collection were rescaled to match the 1.096Å/pix pixel size from published data collections ^100^ using an NRAMM script written for MotionCor2. The CTF of each micrograph was determined using Gctf ^135^. Defocus values for the Retromer/RT-L4 data varied between -0.8 and -2.6 μm; defocus values for the *apo* Retromer data varied between -0.8 and -4.7 μm. RELION-3 ^136^ was used for all image processing unless otherwise indicated.

#### Retromer/RT-L4 processing

Several thousand particles were manually selected to perform initial 2D classification to produce templates for autopicking. Template-based autopicking identified 1,683,975 particles. Multiple rounds of 2D classification yielded 272,349 particles suitable to continue to 3D classification. Initial models for 3D classification were generated by earlier single particle work with wild-type Retromer in the absence of RT-L4 ^100^; models were filtered to 60 Å resolution for use in these experiments. The particles underwent multiple rounds of CTF refinement and Bayesian polishing to produce a final set of 45,330 particles suitable for 3D refinement and postprocessing. The final masked model had a resolution of 5.0 Å and a Relion-determined B-factor of -189.

#### Apo Retromer processing

Data giving rise to the published Retromer *apo* reconstruction lacked tilted views (**Table S4**); an additional data set (**Table S4**; data collection #3) was collected to add tilted views and to improve the reconstruction for this study. Several thousand particles were manually selected from dataset #3 (**Table S4**) to perform initial 2D classification and produce templates for autopicking. Template-based autopicking identified 207,026 particles, which were subjected to initial 2D and 3D classification and refinement as well as CTF refinement. 250,500 particles from data collections #1 and #2 (**Table S4**) Retromer datasets^100^ were imported to combine with data collection #3. Multiple rounds of 2D classification yielded 72,795 particles suitable to continue to 3D classification. Initial models for 3D classification ^100^ were filtered to 60 Å resolution for use in these experiments. The particles underwent multiple rounds of CTF refinement and Bayesian polishing to produce a final set of 43,808 particles suitable for 3D refinement and postprocessing. The final masked model had a resolution of 4.9Å and a Relion-determined B-factor of -113.799.

For both reconstructions, rigid-body docking and map visualization were performed in Chimera^137^ using the Fit in Map routine. Models for N-VPS35 and VPS26A subunits were obtained from PDB 5F0J.

### Liposome preparation

Sucrose-loaded liposome binding assay were performed using the standard extrusion method with some modification ^138^. In brief, cargo loaded Folch liposomes were made by mixing 25 μl of 4 mg/ml N-terminal palmitoylated CIMPR_2347-2376_ peptide, 50 μl of 10 mg/ml Folch fraction I (Sigma Aldrich) and 50 μl of 1 mg/ml di-C16 PtdIns(3)*P* (Echelon Biosciences), each freshly prepared in chloroform, to a total volume of 500 μl chloroform. The solution was dried down on the walls of a mini-round bottom flask under a N_2_ stream and left overnight in a vacuum desiccator to yield a lipid film. This yields liposomes with a final PtdIns(3)P ratio of 10% (w/v). For PC/PE liposome, 50 μl of 10 mg/ml stock of 1-palmitoyl-2-oleoyl-sn-glycero-3-phosphocholine (POPC) and 1-palmitoyl-2-oleoyl phosphatidylethanolamine (POPE) in a 9:1 ratio (Avanti Polar Lipids) was freshly prepared in a 500 μl volume and lipid films formed using the same method as Folch liposomes. This results in a POPC/POPE liposome in a 90%:10% w/v ratio. To form multilamellar vesicles (MLV), the lipid films were hydrated with a buffer comprising 20 mM HEPES pH 7.5 and 220 mM sucrose with agitation follow by10 cycles of rapid freeze-thaw. The sucrose loaded heavy MLVs were centrifuged at 180,000 x *g* for 30 min at 4°C using Optima TL benchtop Ultracentrifuge (Beckman Coulter). The resulting pellets were buffer exchanged by resuspension into the assay buffer comprising 50 mM HEPES pH 7.5, 125 mM NaCl, 0.5 mM TCEP. To avoid buffer mismatch, the protein samples were also buffer exchanged into the same buffer.

### Liposome-Binding Assays

The binding assay was performed in a total volume of 80 µl comprising 40 µl of sucrose-loaded MLVs and 7 µM Retromer, 7 µM SNX3 or 7 µM of Retromer – SNX3 mixture in 1 to 1 ratio. The reaction mixtures were incubated at room temperature for 15 min followed by centrifugation at 36000 x *g* using Optima TL benchtop ultracentrifuge (Beckman Coulter) for 15 min at 4°C. The supernatant and pelleted fractions were then carefully separated. The pellet was then resuspended in 80 µl of buffer containing 50 mM HEPES pH 7.5, 125 mM NaCl, 0.5 mM TCEP before analyzed by SDS-PAGE.

### Statistics

Statistical analysis was completed in R studio using dplyr, ggplot2, ggpubr packages. Error bars on graphs were represented as the standard error of the mean (± SEM). P values were calculated using the two-tailed Student’s *t*-test. P < 0.05 was considered as significant.

### Data deposition

Crystal structural data have been deposited at the Protein data bank (PDB) under the accession number 6XS5 (hVPS29 – RT-D1), 6XS7 (hVPS29 – RT-D2), 6XS8 (ctVPS29 – RT-D3), 6XS9 (hVPS29 – RT-L1), and 6XSA (hVPS29 – RT-L2). CryoEM data has been deposited at the Electron Micrscopy Data Bank (EMDB) under accession numbers D_1000253118 and D_1000253090 for the apo and RT-L4-bound Retromer complexes respectively. All the relevant raw data related to this study is available from the corresponding authors on request.

## Acknowledgments

We acknowledge the use of the Australian Microscopy and Microanalysis Research Facility at the Center for Microscopy and Microanalysis at The University of Queensland. We also acknowledge use of the University of Queensland Remote Operation Crystallization and X-ray (UQ ROCX) Facility and the assistance of Gordon King and Karl Byriel. X-ray data were collected on the MX1 and MX2 microfocus beamline at the Australian Synchrotron. This work is supported by funds from the Australian Research Council (ARC) (DP160101743; DP180103244, CE140100011), National Health and Medical Research Council (NHMRC) (APP1156493; APP1156732), and Bright Focus Foundation (A2018627S). BMC is supported by an NHMRC Senior Research Fellowship (APP1136021), DPF by an NHMRC Senior Principal Research Fellowship (1117017), and DAS is by an NHMRC Career Development Fellowship (APP1140851). This work was also supported by the Japan Agency for Medical Research and Development (AMED), Platform Project for Supporting Drug Discovery and Life Science Research (JP20am0101090) and the Japan Society for the Promotion of Science (JSPS), Specially Promoted Research (JP20H05618) to H.S. CryoEM data were collected at the Vanderbilt Center for Structural Biology Cryo-Electron Microscopy Facility. We thank Dr. Scott Collier and Dr. Melissa Chambers for data collection support. Some of this work was performed at the National Center for CryoEM Access and Training (NCCAT) and the Simons Electron Microscopy Center located at the New York Structural Biology Center, supported by the NIH Common Fund Transformative High Resolution Cryo-Electron Microscopy program (U24 GM129539), and by grants from the Simons Foundation (SF349247) and NY State Assembly. AKK, BX, and LPJ are supported by NIH R35GM119525. LPJ is a Pew Scholar in the Biomedical Sciences, supported by the Pew Charitable Trusts.

## SUPPLEMENTARY INFORMATION

**Figure S1.**
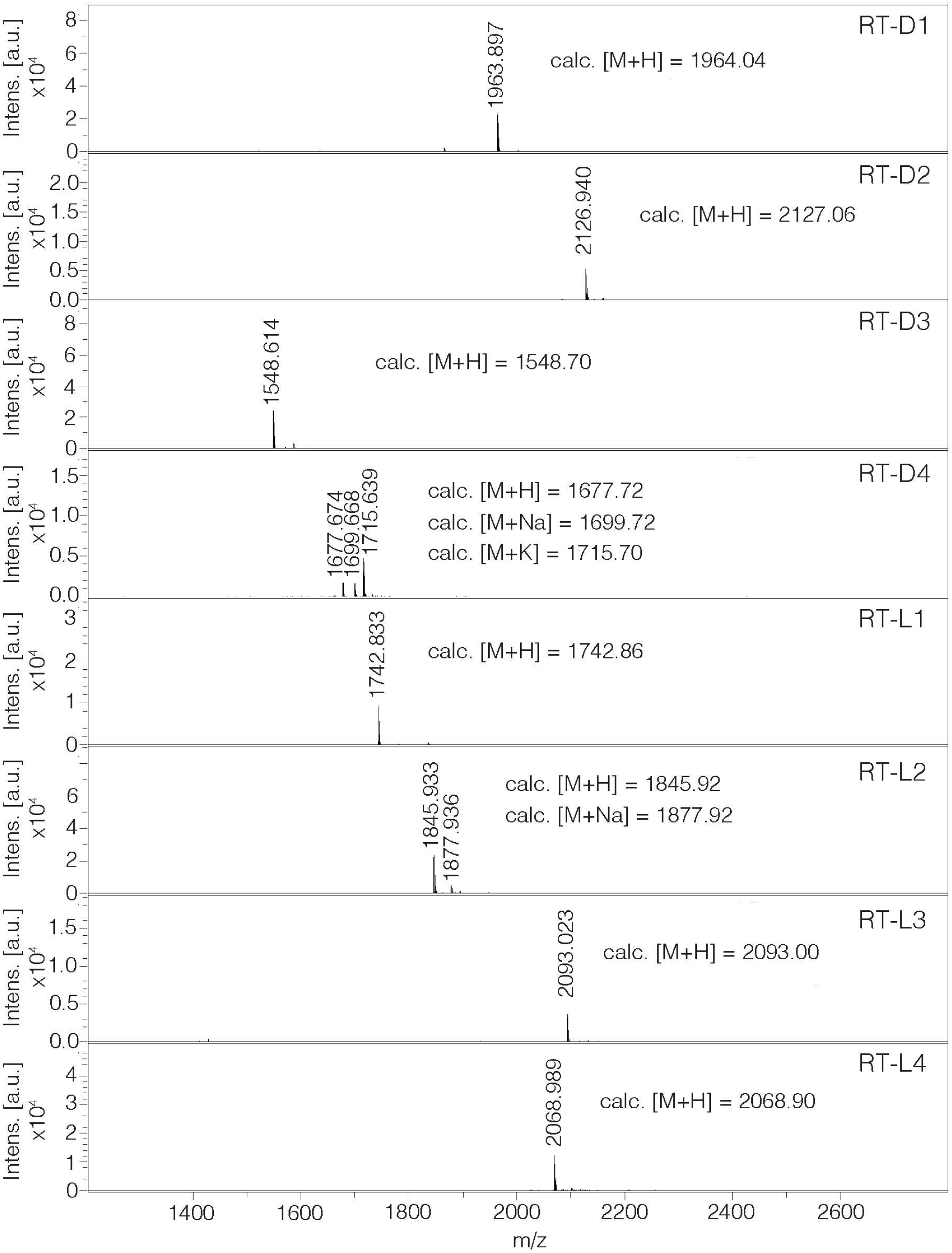
MALDI-TOF spectra of the Retromer associated cyclic peptides. The molecular mass of RT-D1, RT-D2, RT-D3, RT-D4, RT-L1, RT-L2, RT-L3 and RT-L4 are shown in the spectrum.

**Figure S2.**
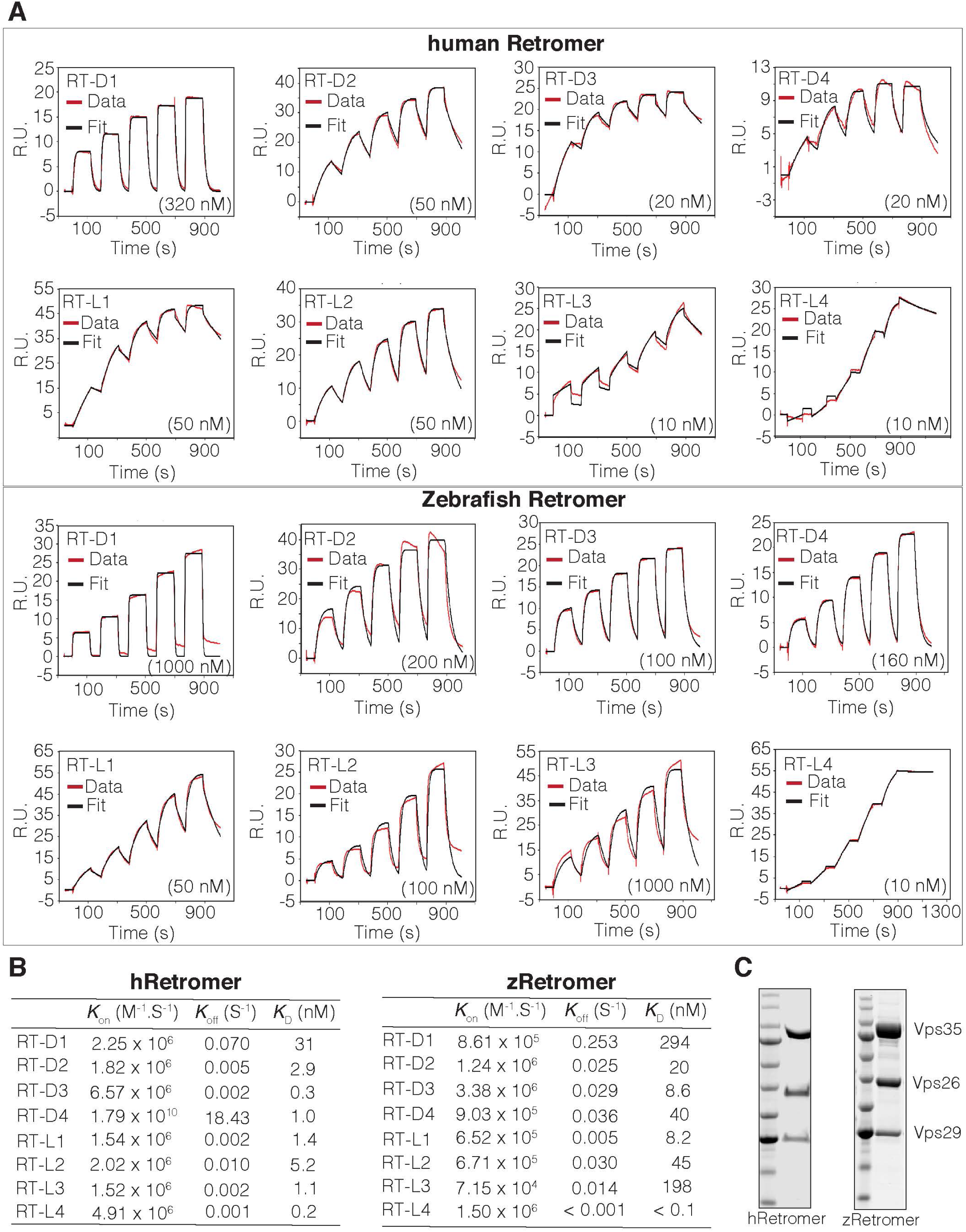
Preliminary SPR binding kinetics of Retromer-associated cyclic peptides. (**A**) Single cycle kinetics experiments were performed using SPR with His-tagged human or zebrafish Retromer with varying concentrations of cyclic peptides. In each case, 2-fold serial dilutions of peptide were tested starting from a highest concentration of 200 or 1000 nM as indicated. (**B**) Binding kinetics of macrocyclic peptides for human and zebrafish Retromer complexes as determined by SPR. (**C**) Gels showing purity of human and zebrafish Retromer complexes used for Rapid peptide screening and SPR experiments.

**Figure S3.**
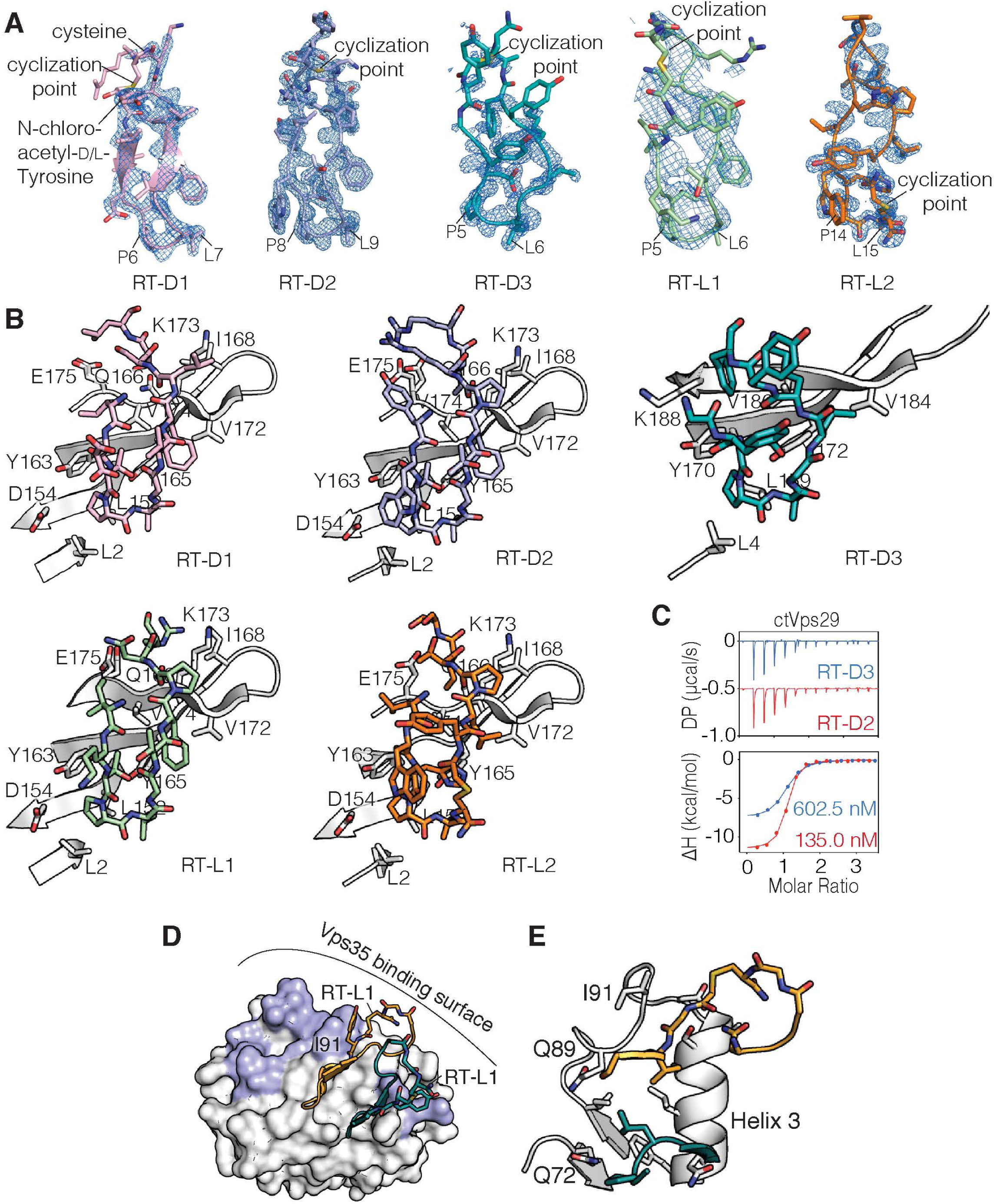
Interaction analysis of Vps29 and cyclic peptides. (**A**) Structures of the Vps29 bound cyclic peptides, RT-D1, RT-D2, RT-D3, RT-L1 and RT-L2. For clarity, the main chain backbones are shown in ribbon and side chains are shown in stick form. The electron density shown corresponds to a simulated-annealing OMIT Fo - Fc map contoured at 3σ. (**B**) Highlighted details of the residues involved in the interactions between Vps29 and bound macrocyclic peptides. (**C**) ITC thermogram for the titration of RT-D2 (red line) and RT-D3 (blue line) with ctVps29. The graph represents the integrated and normalized data fit with a 1 to 1 binding ratio. The binding affinity (*K*_d_) is given as mean of three independent experiments. (**D**) Surface and (**E**) cartoon representations of the human Vps29 – RT-L1 structure highlighting the secondary binding site located opposite to the primary common binding site. Residues involve in contact with Vps35 are shown in light blue.

**Figure S4.**
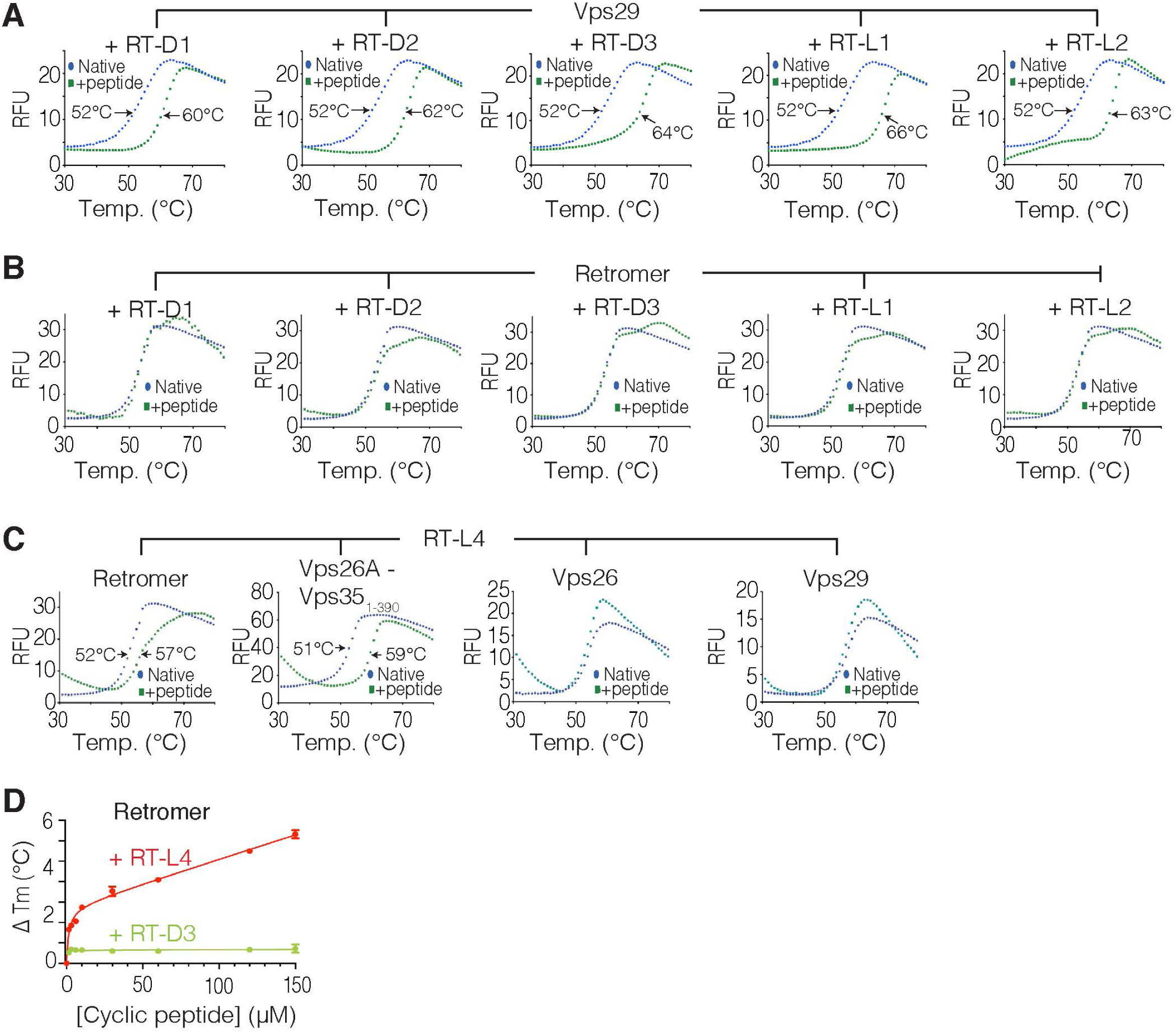
RT-L4 enhances the thermal stability of Retromer in solution. (**A**) Temperature dependent unfolding of Vps29 in the presence of 30-fold molar excess of RT-D1, RT-D2, RT-D3, RT-L1 and RT-L2. (**B**) Temperature dependent unfolding of native Retromer in the presence of 30- fold molar excess of RT-D1, RT-D2, RT-D3, RT-L1 and RT-L2 complexes, and (**C**) in the presence of 30- fold molar excess of RT-L4. Melting temperatures (Tm) were assessed by differential scanning fluorimetry. The sigmoidal curve is characteristic of cooperative thermal denaturation of a folded protein. A shift in melting temperature indicates the stabilization of the proteins upon the addition of cyclic peptide. (**D**) Dose-response curve of Retromer in the presence of RT-L4 (red line) or RT-D3 (green line).

**Figure S5.**
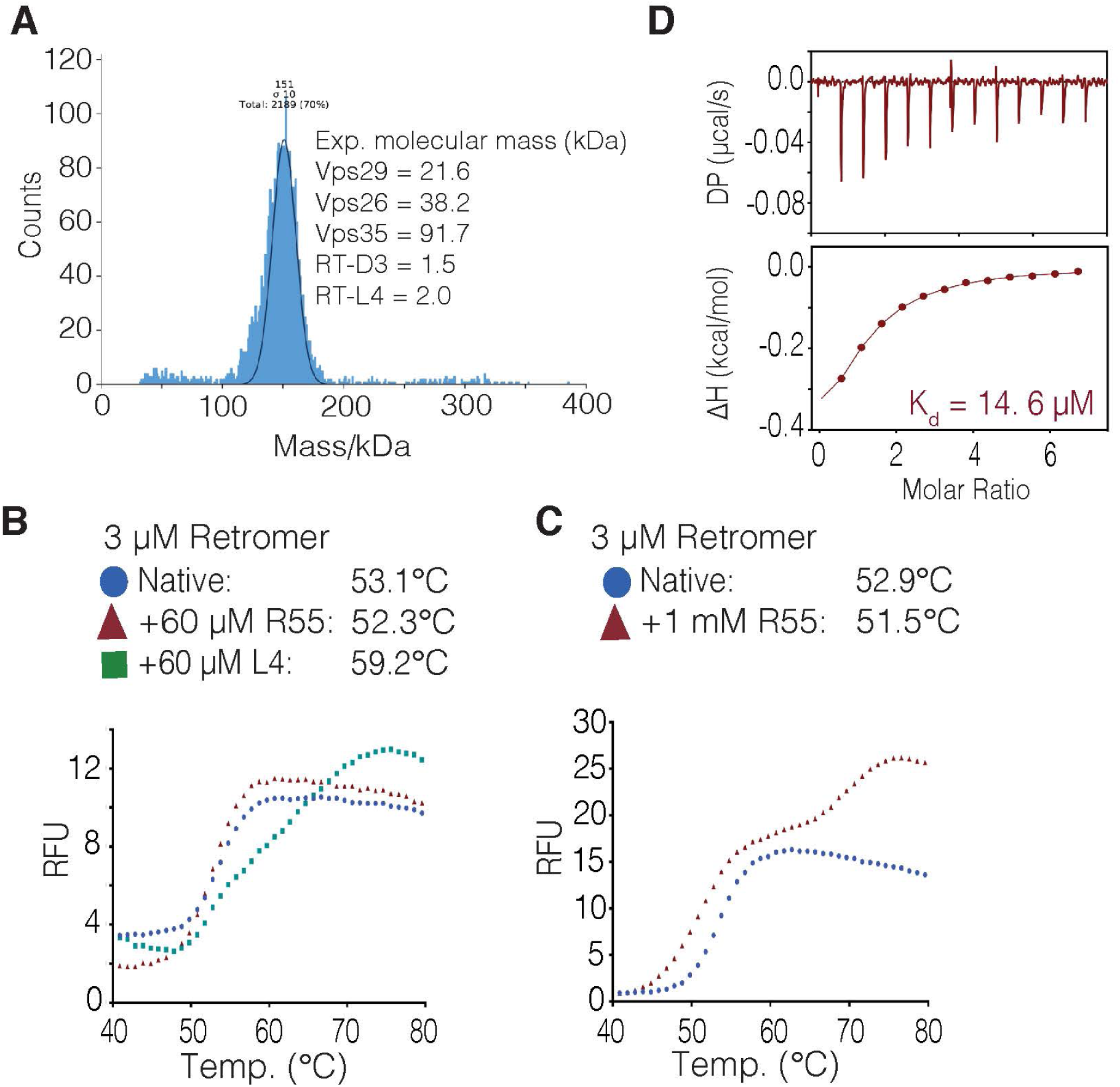
**R55 does not significantly increase the thermal-stability of Retromer. (A**) Molecular mass of Retromer in the presence of RT-D3 and RT-L4 monitored by mass photometry. In this experiment, Retromer shows a mass of 151 kDa, corresponding to the heterotrimeric state of Retromer. (**B**) Temperature dependent unfolding of 3 μM Retromer (blue dot) compared with samples containing 60 μM of R55 (red arrow) or 60 μM RT-L4 (green square). (**C**) Same as (**B**) but with 1 mM of R55 showing two stages of unfolding. Note that preparation of 1 mM RT-L4 in aqueous buffer was not possible due to the lower solubility characteristics. (**D**) ITC thermogram for the titration of R55 with Retromer. The graph represents the integrated and normalized data fit with a 1 to 1 binding ratio. The binding affinity (*K*_d_) is given as mean of three independent experiments.

**Figure S6.**
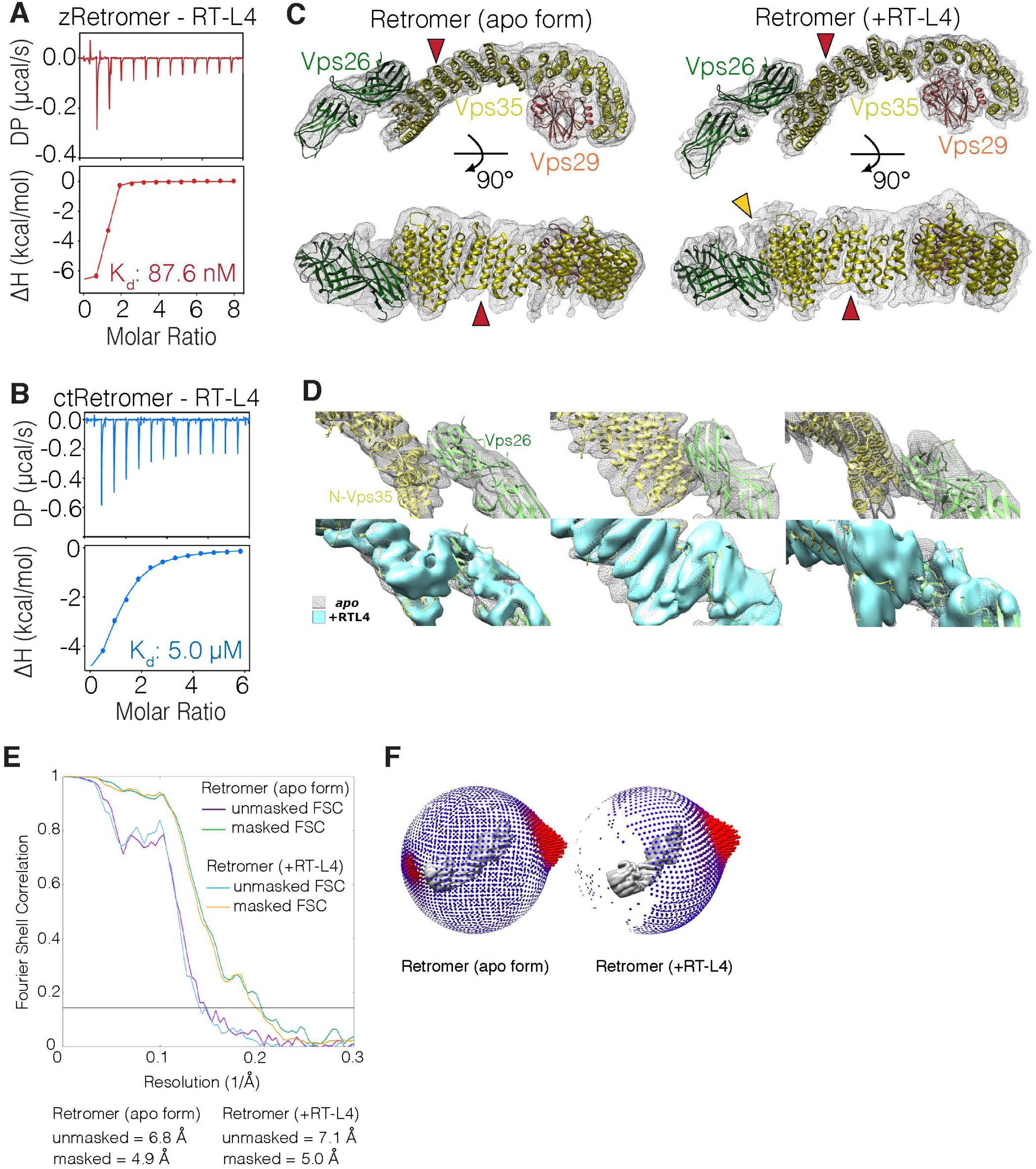
Binding of RT-L4 to Retromer from different species and assessed by cryoEM. (**A**) ITC thermograms for the titration of RT-L4 with zebrafish Retromer showing a strong binding similar to human Retromer. (**B**) ITC thermograms for the titration of RT-L4 with *C. thermophilum* Retromer shows a lower affinity compared to zebrafish or human Retromer. The graphs represents the integrated and normalized data fit with a 1 to 1 binding ratio. The binding affinity (*K*_d_) is given as mean of at least three independent experiments. (**C**) CryoEM density map of human Retromer (apo form) and in complex with RT-L4 displayed as mesh surface, with crystal structures of Retromer subcomplexes (PDB ID 2R17 and 5F0L) overlapped to the map. Red arrow indicates the Vps35 model fitting into α helices. Yellow arrow indicates the extra density observed between Vps26 and Vps35 interface in the RT-L4-bound Retromer. (**D**) Three different views of the Retromer CryoEM structure reconstructions highlighting the Vps35 and Vps26 interface. The cryoEM density from apo Retromer (grey mesh) is overlayed with the cryoEM density from RT-L4-bound Retromer (light blue surface). (**E**) Fourier Shell Correlation (FSC) plots highlighting masked and unmasked resolution estimates from RELION are shown for Retromer (apo form) and in complex with RT-L4. The intersections of the curve with FSC=0.143 (grey dotted line) are shown. (**F**) Angular distribution of the particles used for the final round of refinement. The height and colour of the cylinder bars is proportional to the number of particles in those views.

**Figure S7.**
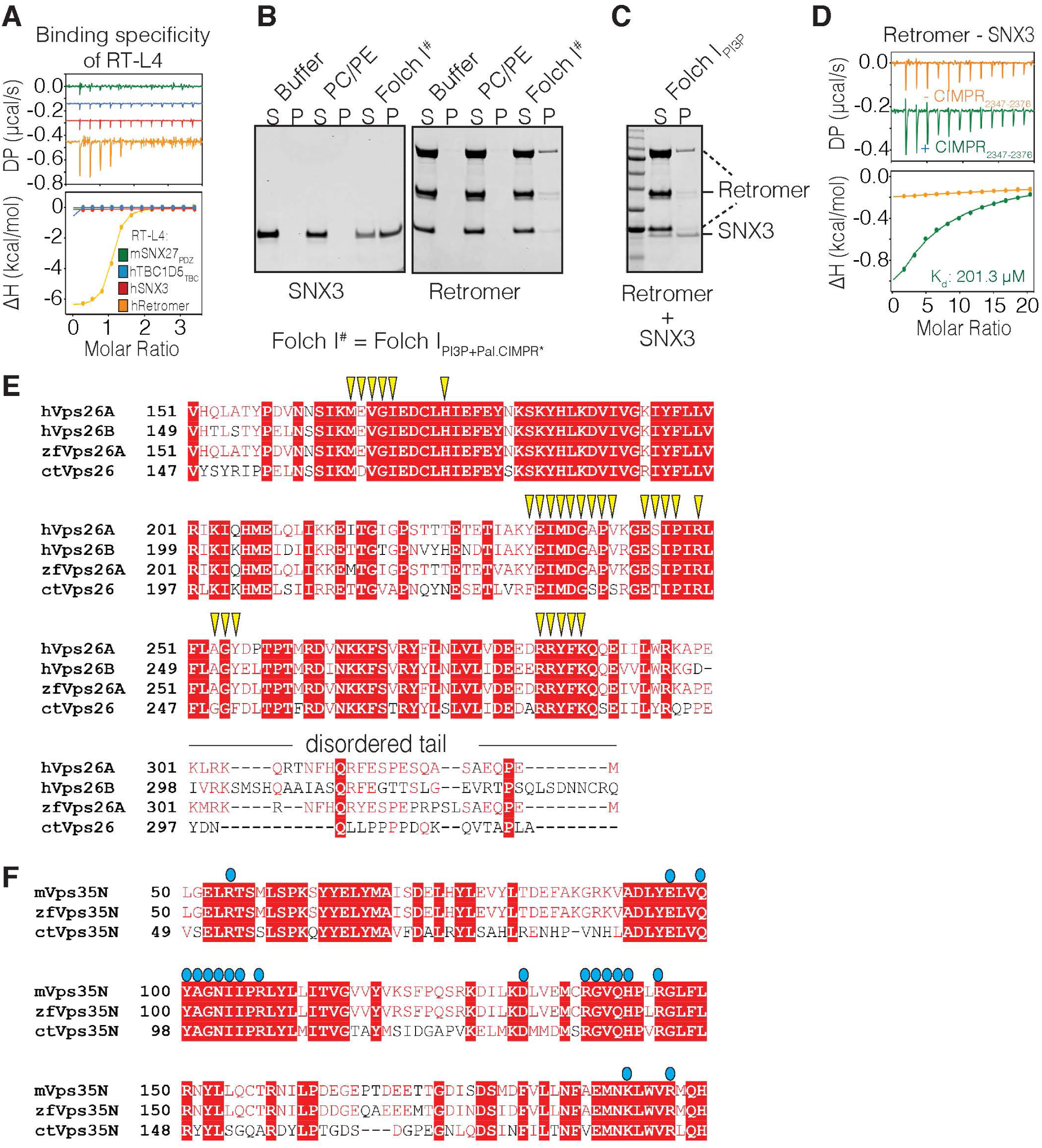
Binding of SNX3 and Retromer in the presence CIMPR_2347-2376_ cargo peptide. (**A**) ITC measurements of RT-L4 with Retromer, SNX27_PDZ_, TBC1D5_TBC_ and SNX3 demonstrate the binding specificity of the cyclic peptide to Retromer. (**B**) Liposome-binding assay of SNX3 and Retromer using Folch I liposomes containing 10% PtdIns(3)*P* and 10% N-terminal palmitoylated CIMPR_2347-2376_ peptide. (**C**) Liposome-binding assay of Retromer and SNX3 mixture as in (**B**) except Folch I liposomes contain only 10% PtdIns(3)*P* without the CI-MPR cargo. Retromer binds only weakly to Folch liposomes in the absence of either SNX3 or cargo sequence. In all three SDS-PAGE gels, “S” indicates unbound supernatant and “P” indicates bound pellet after ultracentrifugation. (**D**) ITC measurement of SNX3 with Vps26A –Vps35_1-390_ subcomplex in the presence of CIMPR_2347-2376_ cargo peptide. The graph represents the integrated and normalized data fit with a 1 to 1 binding ratio. The binding affinity (K_d_) is given as mean of at two three independent experiments. (**E**) Sequence alignment of the C-terminal region of Vps26 and (**F**) the N-terminal region of Vps35 showing the similarity between species. Key residues involve in contacts with Vps35 (yellow arrow) and Vps26 (blue dots) are labelled on top of the sequence. h, *Homo sapiens*; zf, *Danio rerio*; Ct, *Chaetomium thermophilum*; m, *Mus musculus*.

**Figure S8.**
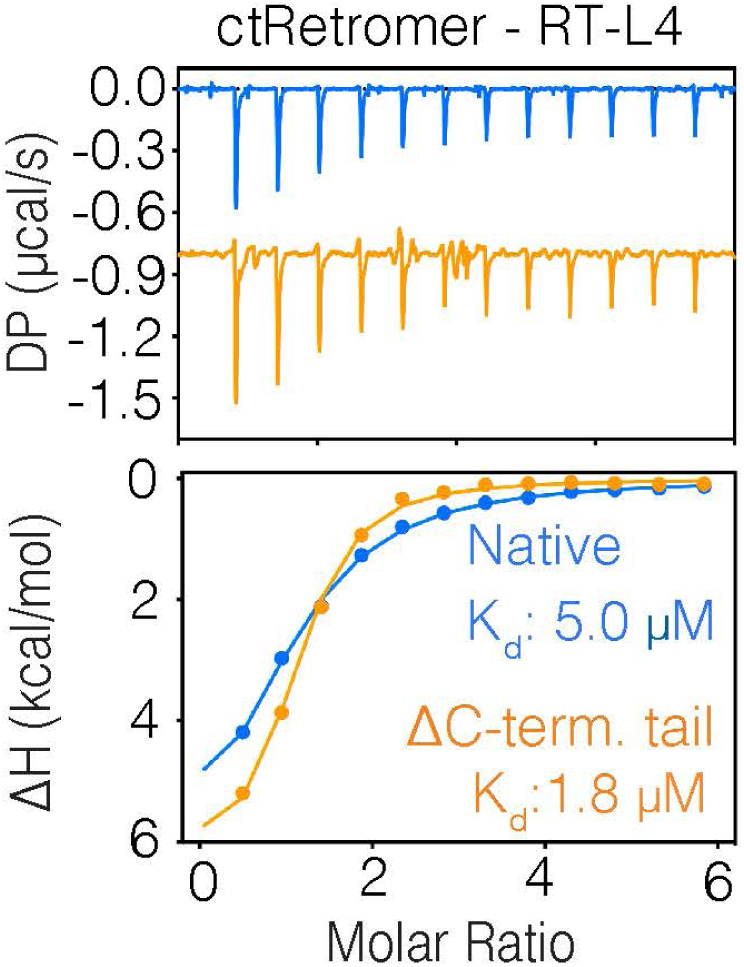
ctVps26 C-terminal disordered tail reveals similar autoinhibitory characteristics. ITC measurement of RT-L4 with native and ctVps26 C-terminal tail truncated ctRetromer. The graph represents the integrated and normalized data fit with a 1 to 1 ratio binding. The binding affinity (K_d_) is given as mean of at least three independent experiments.

**Figure S9.**
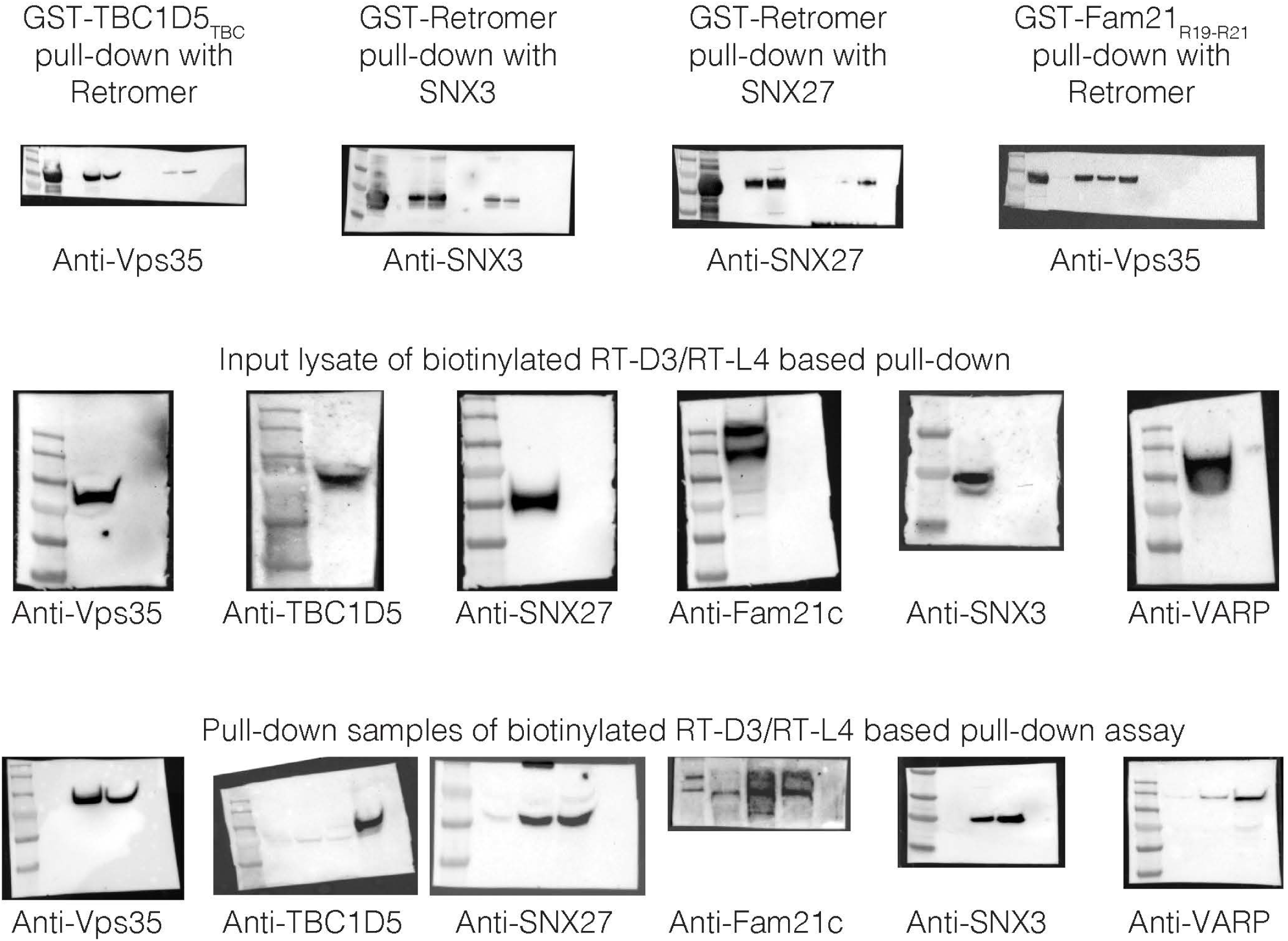
Unprocessed original scans of western blots for the main figure. Unprocessed images of all blots in this study.

**Table S1:**
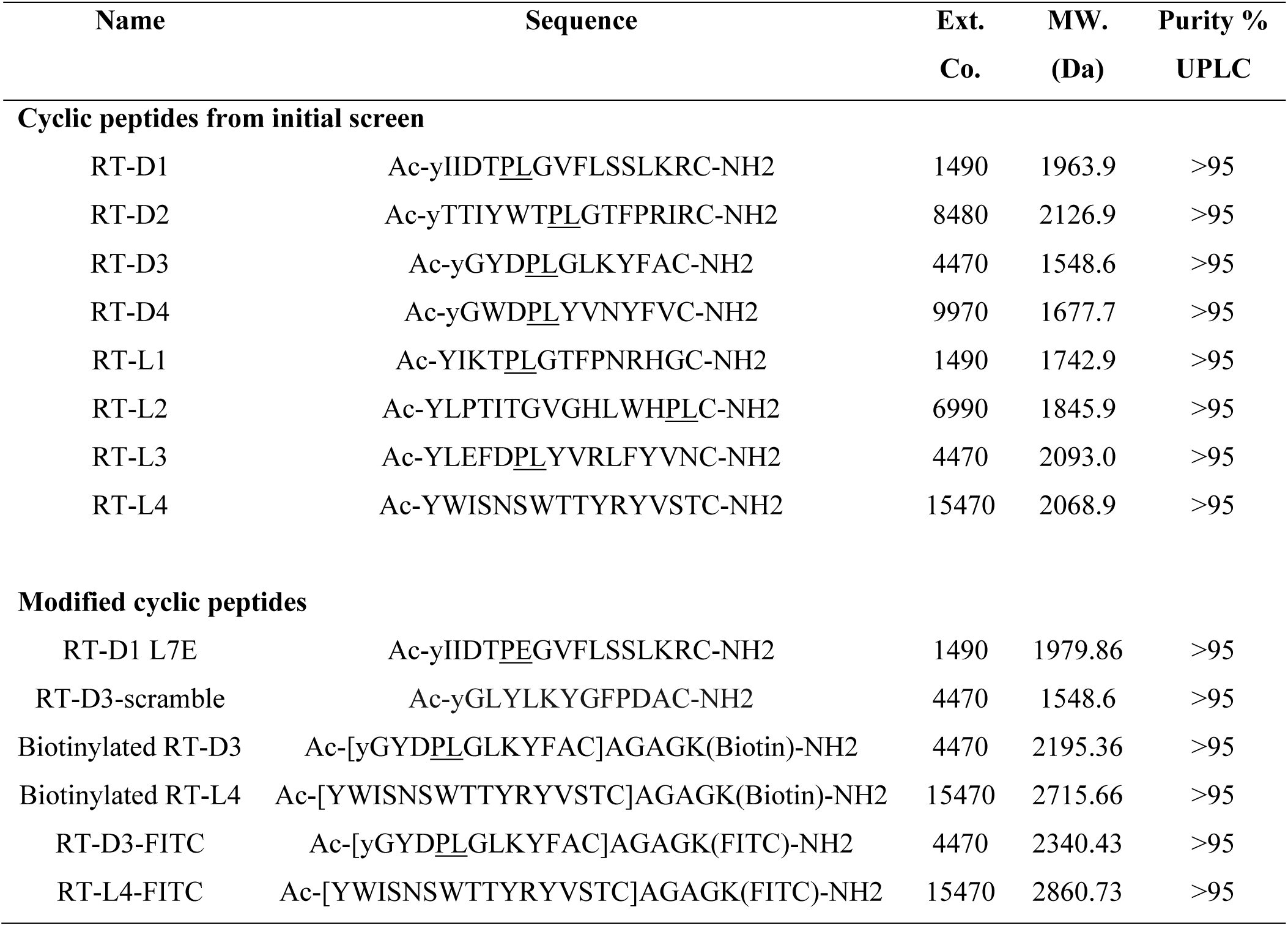
List of synthesized cyclic peptides.

**Table S2:** Thermodynamic parameters for the binding of Retromer with cyclic peptides by ITC.

| | $K_d$<br>(nM) | $\Delta H$<br>(kcal/mol) | $\Delta G$<br>(kcal/mol) | $-T\Delta S$<br>(kcal/mol) |
| --- | --- | --- | --- | --- |
| **Retromer** |  |  |  |  |
| RT-D1 | 130 ± 9 | -11.5 ± 0.5 | -9.4 ± 0.08 | 2.1 ± 0.6 |
| RT-D2 | 53 ± 6 | -9.7 ± 1.6 | -9.9 ± 0.08 | 0.5 ± 0.2 |
| RT-D3 | 25 ± 1 | -13.4 ± 1.4 | -10.4 ± 0.06 | 3.9 ± 0.1 |
| RT-L1 | 96 ± 1 | -11.0 ± 0.9 | -9.5 ± 0.06 | 1.4 ± 0.9 |
| RT-L2 | 852 ± 14 | -16.0 ± 0.7 | -8.2 ± 0.06 | 7.7 ± 0.7 |
| RT-L4 | 273 ± 7 | -7.0 ± 0.1 | -8.9 ± 0.08 | -2.0 ± 0.1 |
| **Vps29** |  |  |  |  |
| RT-D1 | 95 ± 1 | -14.2 ± 0.3 | -9.5 ± 0.06 | 4.6 ± 0.3 |
| RT-D2 | 64 ± 1 | -11.6 ± 0.1 | -9.8 ± 0.08 | 1.7 ± 0.1 |
| RT-D3 | 8 ± 2 | -17.4 ± 1.4 | -11.1 ± 0.21 | 6.4 ± 1.2 |
| RT-L1 | 64 ± 5 | -13.0 ± 1.1 | -9.8 ± 0.08 | 3.2 ± 1.0 |
| RT-L2 | 783 ± 7 | -9.1 ± 1.6 | -8.4 ± 0.06 | 1.6 ± 0.9 |
| RT-L4 |  | No binding detected |  |  |
| **Vps29-Vps35** |  |  |  |  |
| RT-D1 | 106 ± 4 | -10.5 ± 0.4 | -9.5 ± 0.09 | 0.9 ± 0.3 |
| RT-D2 | 65 ± 5 | -11.2 ± 1.0 | -9.8 ± 0.07 | 1.4 ± 0.9 |
| RT-D3 | 29 ± 2 | -11.6 ± 1.3 | -10.3 ± 0.08 | 2.4 ± 1.1 |
| RT-L1 | 90 ± 3 | -7.9 ± 1.2 | -9.6 ± 0.07 | 1.8 ± 1.0 |
| RT-L2 | 763 ± 12 | -8.7 ± 1.7 | -8.4 ± 0.08 | 2.1 ± 1.1 |
| RT-L4 |  | No binding detected |  |  |
| **Vps26-Vps35 truncations** |  |  |  |  |
| Vps26A_Vps35<sub>1-390</sub> : RT-L4 | 165 ± 8 | -6.8 ± 1.2 | -9.2 ± 0.06 | -2.5 ± 1.2 |
| Vps26B_Vps35<sub>1-390</sub> : RT-L4 | 491 ± 27 | -2.7 ± 1.2 | -8.6 ± 0.10 | -5.4 ± 0.4 |
| Vps26A_Vps35<sub>1-224</sub> : RT-L4 | 222 ± 4 | -7.0 ± 1.7 | -9.1 ± 0.11 | -2.1 ± 1.7 |
| Vps26A_Vps35<sub>1-172</sub> : RT-L4 | 339 ± 7 | -6.3 ± 2.5 | -8.8 ± 0.10 | -2.5 ± 2.5 |
| Vps26A<sub>9-298</sub>_Vps35<sub>1-390</sub> : RT-L4<br>(Δ C-term. tail complex) | 33 ± 2 | -11.9 ± 1.0 | -10.2 ± 0.07 | 1.7 ± 1.0 |
| **Vps26A** |  |  |  |  |
| RT-D1, D2, D3, L1, L2, L4 |  | No binding detected |  |  |
| **zfRetromer** |  |  |  |  |
| RT-L4 | 88 ± 11 | -6.9 ± 1.8 | -9.6 ± 0.08 | -2.6 ± 2.0 |
| **ctRetromer** |  |  |  |  |
| RT-L4 | $5030 \pm 900$ | $-7.4 \pm 0.3$ | $-7.3 \pm 0.10$ | $-2.7 \pm 1.9$ |
| **ctRetromer ΔC-term. tail** |  |  |  |  |
| RT-L4 | $1770 \pm 170$ | $-6.0 \pm 0.7$ | $-7.9 \pm 0.06$ | $-1.9 \pm 0.7$ |
| **ctVps29** |  |  |  |  |
| RT-D2 | $135 \pm 3$ | $-10.5 \pm 1.0$ | $-9.3 \pm 0.08$ | $1.1 \pm 0.9$ |
| RT-D3 | $603 \pm 6$ | $-7.4 \pm 0.3$ | $-8.5 \pm 0.07$ | $-0.9 \pm 0.1$ |

**Table S3.**
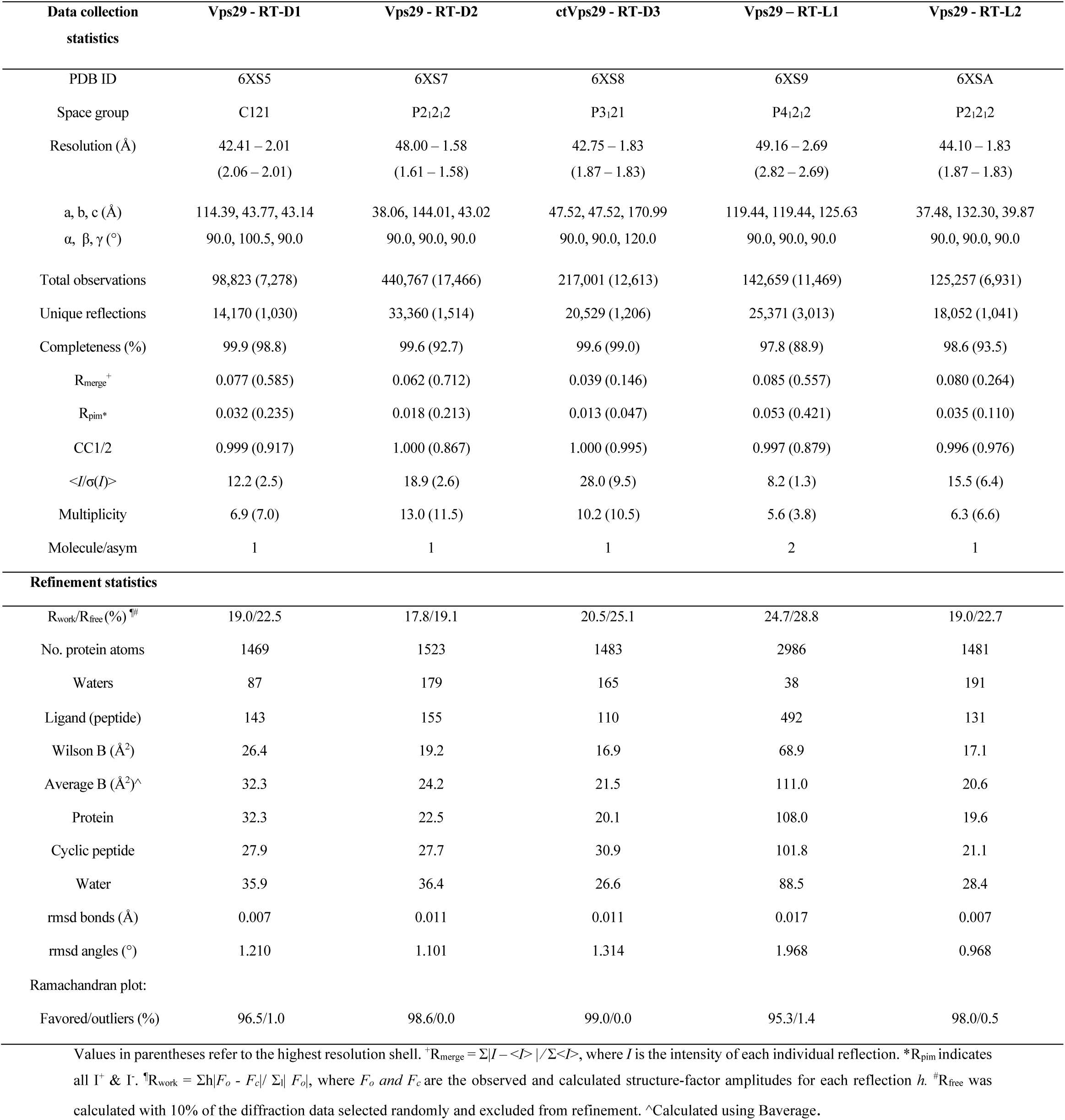
Summary of crystallographic structure determination statistics.

| Data collection statistics | Vps29 - RT-D1 | Vps29 - RT-D2 | ctVps29 - RT-D3 | Vps29 - RT-L1 | Vps29 - RT-L2 |
| --- | --- | --- | --- | --- | --- |
| PDB ID | 6XS5 | 6XS7 | 6XS8 | 6XS9 | 6XSA |
| Space group | C121 | P2<sub>1</sub>2<sub>1</sub>2 | P3<sub>1</sub>21 | P4<sub>1</sub>2<sub>1</sub>2 | P2<sub>1</sub>2<sub>1</sub>2 |
| Resolution (Å) | 42.41 – 2.01<br>(2.06 – 2.01) | 48.00 – 1.58<br>(1.61 – 1.58) | 42.75 – 1.83<br>(1.87 – 1.83) | 49.16 – 2.69<br>(2.82 – 2.69) | 44.10 – 1.83<br>(1.87 – 1.83) |
| a, b, c (Å) | 114.39, 43.77, 43.14 | 38.06, 144.01, 43.02 | 47.52, 47.52, 170.99 | 119.44, 119.44, 125.63 | 37.48, 132.30, 39.87 |
| α, β, γ (°) | 90.0, 100.5, 90.0 | 90.0, 90.0, 90.0 | 90.0, 90.0, 120.0 | 90.0, 90.0, 90.0 | 90.0, 90.0, 90.0 |
| Total observations | 98,823 (7,278) | 440,767 (17,466) | 217,001 (12,613) | 142,659 (11,469) | 125,257 (6,931) |
| Unique reflections | 14,170 (1,030) | 33,360 (1,514) | 20,529 (1,206) | 25,371 (3,013) | 18,052 (1,041) |
| Completeness (%) | 99.9 (98.8) | 99.6 (92.7) | 99.6 (99.0) | 97.8 (88.9) | 98.6 (93.5) |
| R<sub>merge</sub><sup>+</sup> | 0.077 (0.585) | 0.062 (0.712) | 0.039 (0.146) | 0.085 (0.557) | 0.080 (0.264) |
| R<sub>pim</sub><sup>*</sup> | 0.032 (0.235) | 0.018 (0.213) | 0.013 (0.047) | 0.053 (0.421) | 0.035 (0.110) |
| CC1/2 | 0.999 (0.917) | 1.000 (0.867) | 1.000 (0.995) | 0.997 (0.879) | 0.996 (0.976) |
| <I/σ(I)> | 12.2 (2.5) | 18.9 (2.6) | 28.0 (9.5) | 8.2 (1.3) | 15.5 (6.4) |
| Multiplicity | 6.9 (7.0) | 13.0 (11.5) | 10.2 (10.5) | 5.6 (3.8) | 6.3 (6.6) |
| Molecule/asym | 1 | 1 | 1 | 2 | 1 |
| **Refinement statistics** |  |  |  |  |  |
| R<sub>work</sub>/R<sub>free</sub> (%) <sup>#</sup> | 19.0/22.5 | 17.8/19.1 | 20.5/25.1 | 24.7/28.8 | 19.0/22.7 |
| No. protein atoms | 1469 | 1523 | 1483 | 2986 | 1481 |
| Waters | 87 | 179 | 165 | 38 | 191 |
| Ligand (peptide) | 143 | 155 | 110 | 492 | 131 |
| Wilson B (Å<sup>2</sup>) | 26.4 | 19.2 | 16.9 | 68.9 | 17.1 |
| Average B (Å<sup>2</sup>)<sup>^</sup> | 32.3 | 24.2 | 21.5 | 111.0 | 20.6 |
| Protein | 32.3 | 22.5 | 20.1 | 108.0 | 19.6 |
| Cyclic peptide | 27.9 | 27.7 | 30.9 | 101.8 | 21.1 |
| Water | 35.9 | 36.4 | 26.6 | 88.5 | 28.4 |
| rmsd bonds (Å) | 0.007 | 0.011 | 0.011 | 0.017 | 0.007 |
| rmsd angles (°) | 1.210 | 1.101 | 1.314 | 1.968 | 0.968 |
| Ramachandran plot: |  |  |  |  |  |
| Favored/outliers (%) | 96.5/1.0 | 98.6/0.0 | 99.0/0.0 | 95.3/1.4 | 98.0/0.5 |

Values in parentheses refer to the highest resolution shell. <sup>+</sup>R<sub>merge</sub> = Σ|I – <I>| / Σ<I>, where I is the intensity of each individual reflection. <sup>\*</sup>R<sub>pim</sub> indicates all I<sup>+</sup> & I<sup>-</sup>. <sup>#</sup>R<sub>work</sub> = Σh|F<sub>o</sub> – F<sub>c</sub>| / Σ|F<sub>o</sub>|, where F<sub>o</sub> and F<sub>c</sub> are the observed and calculated structure-factor amplitudes for each reflection h. <sup>#</sup>R<sub>free</sub> was calculated with 10% of the diffraction data selected randomly and excluded from refinement. <sup>^</sup>Calculated using Baverage.

**Table S4:** CryoEM data collection parameters for 3KE Retromer complex in the presence of the RT-L4 cyclic peptide.

|  | **Retromer + RT-L4** |  | ***apo* Retromer** |  |
| --- | --- | --- | --- | --- |
| **EMDB ID** | D_1000253090 |  | D_1000253118 |  |
| **Microscope** | ThermoFisher FEI Titan Krios G3i Vanderbilt V-CEM | FEI Titan Krios Krios2 NRAMM Data collection 1 | FEI Titan Krios Krios3 NRAMM Data collection 2 | FEI Titan Krios Krios3 NRAMM Data collection 3 |
| **Cs** | 2.7 mm | 2.7 mm | 2.7 mm | 2.7 mm |
| **Voltage** | 300 keV | 300 keV | 300 keV | 300 keV |
| **Detector** | ThermoFisher Falcon 3 direct electron detector | Gatan K2 Summit direct electron detector | Gatan K2 Summit direct electron detector | Gatan K2 Summit direct electron detector |
| **Magnification** | 120,000x | 105,000x | 105,000x | 105,000x |
| **Pixel size** | 0.6811Å/pix | 1.096Å/pix | 1.06Å/pix (rescaled to 1.096Å/pix with motioncorr2 script) | 1.06Å/pix (rescaled to 1.096Å/pix with motioncorr2 script) |
| **Dose rate** | 1.4e<sup>-</sup>/Å<sup>2</sup>/sec | ~8e<sup>-</sup>/Å<sup>2</sup>/sec | ~8e<sup>-</sup>/Å<sup>2</sup>/sec | ~8e<sup>-</sup>/Å<sup>2</sup>/sec |
| **Total dose** | 50e<sup>-</sup>/Å<sup>2</sup> | 69.34e<sup>-</sup>/Å<sup>2</sup> | 73.92e<sup>-</sup>/Å<sup>2</sup> | 73.92e<sup>-</sup>/Å<sup>2</sup> |
| **Tilt** | +/-30° | N/A | N/A | +/-15° |
| **Defocus range** | -0.8 to -2.6µm | -0.7 to 2.6µm | -0.8 to -4.4µm | -0.8 to -4.7µm |
| **Number of micrographs** | 4,791 | 1,480 | 1,299 | 891 |
| **Total particles (autopicked)** | 1,683,975 | (250,500 particles selected from these datasets; Kendall *et al.*, 2020) |  | 207,026 |
| **Box size (Å)** | 501 Å |  | 241 Å |  |
| **Particles in 2D classification** | 273,172 |  | 72,795 |  |
| **Particles in final 3D model** | 45,330 |  | 43,808 |  |
| **Symmetry** | C1 |  | C1 |  |
| **Map resolution (masked FSC 0.143, RELION)** | 5.0 Å |  | 4.9Å |  |
| **B-factor** | -189 |  | -114 |  |

**Table S5:** Thermodynamic parameters for the binding of Retromer and its partners in the presence of RT-D3 or RT-L4 cyclic peptides.

| | $K_d$<br>($\mu\text{M}$) | $\Delta H$<br>(kcal/mol) | $\Delta G$<br>(kcal/mol) | $-T\Delta S$<br>(kcal/mol) |
| --- | --- | --- | --- | --- |
| **TBC1D5<sub>TBC</sub>** |  |  |  |  |
| Retromer + DMSO (Native) | $0.37 \pm 0.02$ | $-2.5 \pm 0.8$ | $-8.8 \pm 0.03$ | $-6.2 \pm 0.8$ |
| Retromer + RT-D3 |  | No binding detected |  |  |
| Retromer + RT-L4 | $0.19 \pm 0.01$ | $-5.7 \pm 0.1$ | $-9.2 \pm 0.06$ | $-3.5 \pm 0.1$ |
| **SNX27<sub>PDZ</sub>** |  |  |  |  |
| Retromer + DMSO (Native) | $13.0 \pm 0.5$ | $-1.1 \pm 0.1$ | $-6.7 \pm 0.03$ | $-5.6 \pm 0.2$ |
| Retromer + RT-D3 | $15.1 \pm 0.4$ | $-1.2 \pm 0.3$ | $-6.6 \pm 0.08$ | $-5.4 \pm 0.2$ |
| Retromer + RT-L4 | $8.2 \pm 1.0$ | $-5.5 \pm 0.3$ | $-6.9 \pm 0.07$ | $-1.4 \pm 0.4$ |
| **PTHR<sub>586-593</sub>** |  |  |  |  |
| mSNX27<sub>PDZ</sub> | $5.2 \pm 0.4$ | $-10.6 \pm 0.4$ | $-7.2 \pm 0.05$ | $3.3 \pm 0.4$ |
| Retromer + mSNX27<sub>PDZ</sub> | $2.4 \pm 0.3$ | $-11.1 \pm 1.8$ | $-7.7 \pm 0.08$ | $3.5 \pm 1.9$ |
| Retromer + mSNX27<sub>PDZ</sub> + RT-L4 | $0.7 \pm 0.1$ | $-12.5 \pm 0.6$ | $-8.4 \pm 0.05$ | $4.3 \pm 0.3$ |
| **SNX3** |  |  |  |  |
| Retromer + DMT1* + DMSO (Native) | $154 \pm 25$ | $-32.4 \pm 9.8$ | $-5.2 \pm 0.14$ | $27.1 \pm 9.8$ |
| Retromer + DMT1* + RT-D3 | $164 \pm 3$ | $-33.2 \pm 0.4$ | $-5.2 \pm 0.02$ | $28.0 \pm 0.4$ |
| Retromer + DMT1* + RT-L4 | $230.0 \pm 14$ | $-14.0 \pm 0.6$ | $-5.0 \pm 0.03$ | $9.0 \pm 0.6$ |
| **GST-Fam21<sub>R19-R21</sub>** |  |  |  |  |
| Retromer + DMSO (Native) | $10.6 \pm 0.7$ | $-16.5 \pm 0.6$ | $-6.8 \pm 0.04$ | $9.8 \pm 0.9$ |
| Retromer + RT-D3 | $25.8 \pm 0.5$ | $-9.3 \pm 3.3$ | $-6.3 \pm 0.02$ | $4.0 \pm 1.9$ |
| Retromer + RT-L4 | $2.8 \pm 0.8$ | $-26.0 \pm 1.6$ | $-7.5 \pm 0.02$ | $18.4 \pm 1.8$ |

DMT1\* corresponds to DMT1-II<sub>550-568</sub>.

**Table S6.** Thermodynamic parameters for the binding of Vps26 – Vps35 subcomplex with SNX3 in the presence of cargo peptide.

| | $K_d$<br>(μM) | $\Delta H$<br>(kcal/mol) | $\Delta G$<br>(kcal/mol) | $-T\Delta S$<br>(kcal/mol) |
| --- | --- | --- | --- | --- |
| **Vps26A – Vps35<sub>1-390</sub> – DMT1-II<sub>550-568</sub> : SNX3** |  |  |  |  |
| Native | 149 ± 9 | -14.6 ± 0.3 | -5.2 ± 0.05 | 10.4 ± 1.6 |
| Phosphomimetic mutant (pm) | 61 ± 3 | -18.7 ± 0.7 | -5.8 ± 0.03 | 13.0 ± 0.8 |
| QRFE to AAAA mutant | 50 ± 2 | -23.0 ± 3.2 | -5.9 ± 0.03 | 17.1 ± 3.0 |
| ΔC-term. tail (ΔC) | 32 ± 1 | -28.3 ± 0.2 | -6.1 ± 0.04 | 22.1 ± 0.4 |
| **Vps26A – Vps35<sub>1-390</sub> – CIMPR<sub>2347-2376</sub> : SNX3** |  |  |  |  |
| Native | 201 ± 9 | -11.8 ± 0.9 | -5.1 ± 0.03 | 6.6 ± 0.8 |
| **Vps26A – Vps35<sub>1-390</sub> : SNX3** |  |  |  |  |
| Native |  | No binding detected |  |  |
| Phosphomimetic mutant (pm) |  | No binding detected |  |  |
| ΔC-term.tail (ΔC) | | No binding detected | | |
| **Vps26B – Vps35<sub>1-390</sub> – DMT1-II<sub>550-568</sub> : SNX3** |  |  |  |  |
| Native | 193 ± 4 | -13.8 ± 0.5 | -5.1 ± 0.02 | 8.7 ± 0.6 |
| Phosphomimetic mutant (pm) | 64 ± 2 | -12.3 ± 2.3 | -5.7 ± 0.02 | 7.1 ± 1.5 |
| ΔC-term.tail (ΔC) | 34 ± 2 | -24.7 ± 2.7 | -6.1 ± 0.03 | 18.6 ± 2.7 |
| **Vps26B – Vps35<sub>1-390</sub> : SNX3** |  |  |  |  |
| Native |  | No binding detected |  |  |
| Phosphomimetic mutant (pm) |  | No binding detected |  |  |
| ΔC-term.tail (ΔC) | | No binding detected | | |

